# Excitatory-inhibitory balance modulates the formation and dynamics of neuronal assemblies in cortical networks

**DOI:** 10.1101/2021.04.15.439946

**Authors:** Sadra Sadeh, Claudia Clopath

## Abstract

Repetitive activation of subpopulation of neurons in cortical networks leads to the formation of neuronal assemblies, which can guide learning and behavior. Recent technological advances have made the artificial induction of such assemblies feasible, yet how various patterns of activation can shape their emergence in different operating regimes is not clear. Here we studied this question in large-scale cortical networks composed of excitatory (E) and inhibitory (I) neurons. We found that the dynamics of the network in which neuronal assemblies are embedded is important for their induction. In networks with strong E-E coupling at the border of E-I balance, increasing the number of perturbed neurons enhanced the potentiation of ensembles. This was, however, accompanied by off-target potentiation of connections from unperturbed neurons. When strong E-E connectivity was combined with dominant E-I interactions, formation of ensembles became specific. Counter-intuitively, increasing the number of perturbed neurons in this regime decreased the average potentiation of individual synapses, leading to an optimal assembly formation at intermediate sizes. This was due to potent lateral inhibition in this regime, which also slowed down the formation of neuronal assemblies, resulting in a speed-accuracy trade-off in the performance of the networks in pattern completion and behavioral discrimination. Our results therefore suggest that the two regimes might be suited for different cognitive tasks, with fast regimes enabling crude detections and slow but specific regimes favoring finer discriminations. Functional connectivity inferred from recent experiments in mouse cortical networks seems to be consistent with the latter regime, but we show that recurrent and top-down mechanisms can dynamically modulate the networks to switch between different states. Our work provides a framework to study how neuronal perturbations can lead to network-wide plasticity under biologically realistic conditions, and sheds light on the design of future experiments to optimally induce behaviorally relevant neuronal assemblies.

## Introduction

Neuronal assemblies are building blocks of computation and learning in the brain (*1–4*). Recent technological advances have provided us with unprecedented tools to bidirectionally interact with their circuitry, by enabling us to record from and perturb the activity of subpopulations of neurons (*5–7*), in order to ultimately link their dynamics to behavior (*8*). Experimental work has specially been successful in recent years to artificially induce neuronal ensembles by targeted activation of a subset of cortical neurons (*9, 10*). Efficient induction of such assemblies can provide a powerful means to trigger or suppress a specific behavior (*8, 11*), and can potentially guide us in understanding brain diseases (*12*) and to design more efficient brain machine interfaces (*13*). As with other perturbation techniques (*14–17*), it is crucial to understand how parameters of stimulation, including the pattern of activation of specific neurons and the general state of the network dynamics, can be optimized for an efficient induction.

This optimization can, however, be complicated, given the complex connectivity and dynamics of cortical networks. Neuronal ensembles cannot form in isolation, as they are embedded in a background of connections from other neurons in local and distal networks, which can modulate their activity. The background network itself can be in different operating regimes (e.g. awake versus anaesthetized) and modulated by different mechanisms like top-down input and neuromodulators. Moreover, perturbation of a subset of neurons embedded in such background networks is likely to create a cascade of activation of downstream neurons (*18, 19*), which can in turn affect the activity and plasticity of the initially perturbed neurons in a recurrent manner. This complex interaction is especially expected in cortical networks with strong recurrent excitatory and inhibitory connections, as reported in many brain regions (*19–25*).

To study the formation of neuronal assemblies under biologically realistic conditions we therefore need to consider this complex interplay of network dynamics and plasticity. Conventional plasticity protocols, in contrast, typically study the weight changes under artificial conditions (*26–28*). The plasticity is often induced in a few, isolated pairs of neurons, in conditions where the effect of background activity (*29*) and network interactions is masked or minimized. They also involve patterns (*30*) and conditions (*31*) of stimulation that are optimized to maximally drive the activity of pre- and post-synaptic neurons (**Fig. 1A**). Learning in naturalistic conditions, on the other hand, is likely to be guided by a different pattern of neuronal perturbations (*30*), involving sparse activity of a large subpopulation of neurons (*32*), spanning a wide range of spatial and temporal scales pertinent to behaviourally relevant stimuli (*33*) (**Fig. 1B**).

**Fig. 1.**
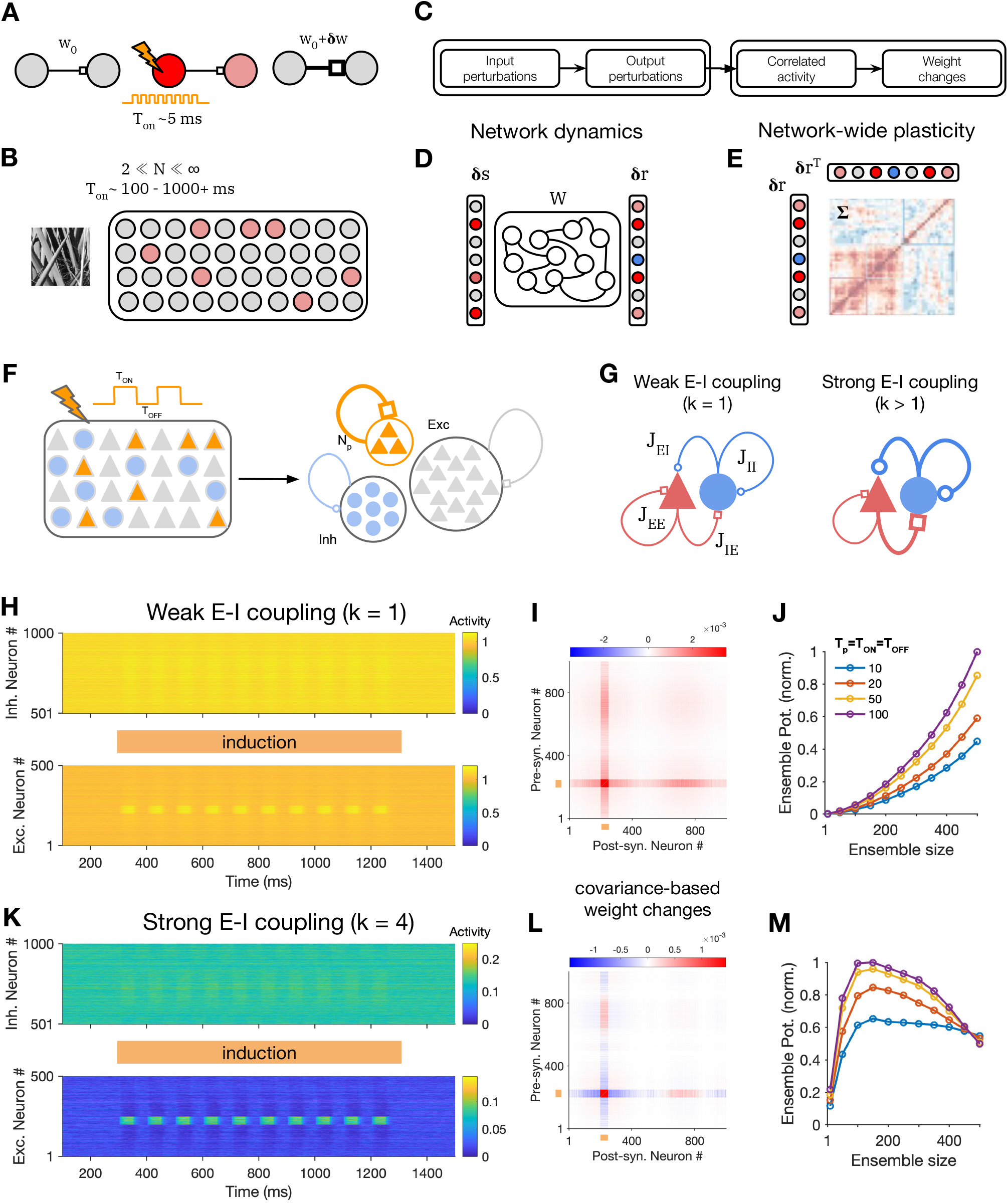
Induction of neuronal assemblies in different regimes of excitation-inhibition balance. (**A**) Schematic of conventional protocols for the induction and investigation of plasticity, often involving a small number of neurons and perturbations with brief ON pulses (*T*_*on*_). (**B**) Typical activity patterns in response to naturalistic stimuli involves activation of a large but sparsely active subset of neurons, with typical temporal scales much longer than a few milliseconds (*33*). (**C-E**) Analytical steps (C) to evaluate the effect of external perturbations on the formation of neuronal assemblies, involving dynamics of networks responses (D) and network-wide plasticity (E). Knowing the weight matrix (*W*), input perturbations (δ*s*) are transferred to output perturbations (*δr*) (D); the resulting correlated activity patterns of pre- and post-synaptic neurons (Σ) in turn guide a network-wide plasticity of weights (E). (**F**) Schematic of the perturbation protocol to study plasticity in large-scale networks, composed of excitatory (Exc, triangles) and inhibitory (Inh, circles) neurons. Left: *N*_*p*_ excitatory neurons are perturbed, with a series of stimulation pulses which are alternating between on and off states for *T*_*on*_ and *T*_*off*_, respectively. Right: The effect of perturbations on the induction of assemblies is assayed by evaluating the potentiation of weights within the perturbed neurons (orange) as a result of Hebbian learning (see **Methods**). (**G**) Parameterization of different regimes of large-scale networks in which neuronal assemblies are induced, with two sample networks demonstrating weak (*k* = 1) and strong (*k* > 1) E-I coupling regimes. *J*_IE_ = |*J*_IE_| = |*J*_II_| = *kJ*_*EE*_. (**H**) Responses of *N*_E_ excitatory and *N*_I_ inhibitory neurons in a network with weak E-I coupling (*k* = 1) to perturbations (10 pulses with *T*_*p*_ = 50, delivered to *N*_*p*_ = 50 neurons, starting from *T* = 300). *N*_E_ = *N*_I_ = 500. (**I**) Matrix of weight changes resulting from a Hebbian plasticity rule based on the covariance of response changes after perturbations in (H). Neurons are sorted such that closer neurons are more strongly connected in the initial weight matrix (see **Methods** for details). Orange bars denote the stimulated neurons. (**J**) Ensemble potentiation for different ensemble sizes (*N*_*p*_) resulting from perturbations with different temporal profiles (*T*_*p*_). Ensemble potentiation is evaluated, within the assembly of perturbed neurons, as the normalized average sum of all pre-synaptic weights to perturbed neurons (averaged across post-synaptic sources and normalized to the maximum ensemble potentiation for tested combinations of *N*_*p*_ and *T*_*p*_). (**K-M**) Same as (H-J) for networks with strong E-I coupling (*k* = 4).

To breach this gap, here we studied how neuronal ensembles can be induced in large-scale recurrent networks of excitatory and inhibitory neurons under various regimes of perturbation. We used a theory we recently developed to analyse the effect of neuronal perturbations in such networks (*34*), and asked how activity changes resulting from different perturbations guide network-wide plasticity (**Fig. 1C-E**). Network response dynamics operate on much faster timescales (∼10-100 ms) compared to the timescale of long-term plasticity (minutes to days), so we used separation of timescales between the two stages to guide our analysis. In the first step, we assessed how input perturbations are transferred to output responses, assuming a fixed weight matrix for the network (**Fig. 1D**); we then studied how the correlative activity patterns emerging from such responses lead to modification of neuronal ensembles (**Fig. 1E**). We started our study by analyzing the influence of network dynamics on network-wide plasticity (**Fig. 1C**), but later also considered the bidirectional interaction of the two over longer time scales.

Using this approach we found that not only the parameters of external stimulation, but also the operating regime of the network affect the induction of neuronal assemblies. Ensembles formed fast but with less specificity in networks with weak E-I coupling, while networks with strong E-I coupling gave rise to specific ensembles, which formed more slowly. Moreover, the size of perturbed neurons had different effects on the formation of assemblies in different regimes. Specifically, increasing the number of perturbed neurons in strong E-I coupling regimes did not always enhance the formation of assemblies, suggesting an optimal size for the induction of ensembles. Different E-I regimes also favored a different trade-off of speed and accuracy in behavioral tasks, and modulating the E-I regime provided a powerful and generic means to control this trade-off in assembly formation and learning.

## Results

### Induction of neuronal assemblies in excitatory-inhibitory networks

We studied the formation of neuronal assemblies as a result of different patterns of perturbations in large-scale cortical network models with balance of excitation (E) and inhibition (I) (*35–37*). A subset of excitatory neurons is targeted by repetitive external perturbations to induce a neuronal assembly (**Fig. 1F**). Induction protocols are characterized by the key parameters of the perturbations including the number of targeted neurons (*N*_*p*_ – or the size of neuronal ensemble), and the temporal properties of the stimulus, which is alternating between binary states of ON (*S* = 1, for *T*_*ON*_) and OFF (*S* = 0, for *T*_*OFF*_). The background network is parameterized by the strength of E-E weights (*J*_*EE*_ = *J*), and the relative strength of E-I coupling (*k*). The parameter *k* describes the dominance of the overall *E* → *I* and *I* → {E, I} couplings relative to E-E weights, with *k* = 1 denoting perfect balance and *k* > 1 ensuring a dominant recurrent inhibition (**Fig. 1G**; see **Methods**). E-E coupling is strong (*J* > 1), such that in the absence of E-I interactions (*k* = 0) the network is unstable, consistent with the observation that cortical networks operate in inhibitory-stabilized regimes of activity (*24, 38*).

We simulated the response of the network to perturbations in two regimes of recurrent E-I interaction (**Fig. 1H-M**). The first regime is equipped with the minimum amount of inhibition that is necessary to stabilize the E-E subnetwork (*k* = 1); we refer to this regime as weak E-I coupling (**Fig. 1G**, left). The second regime has a stronger E-I coupling (*k* = 4) (**Fig. 1G**, right), which guarantees the operation of the network away from the border of instability and enables stronger lateral inhibition. We simulate the response of the network before and after perturbations in each regime (**Fig. 1H,K**), and evaluate how Hebbian-type plasticity rules (see **Methods**) change the weight matrix as a result of pre- and post-synaptic activity changes (**Fig. 1I,L**). The strength of neuronal ensemble resulting from perturbation-induced plasticity was quantified by calculating the potentiation of synapses within the perturbed subset of neurons in each case (**Fig. 1J,M**).

Networks with weak E-I balance (*k* = 1) showed supralinear potentiation of the strength of assemblies with increasing the size of targeted neurons (*N*_*p*_), and this effect was increased for longer time interval of perturbation (*T*_*p*_) (**Fig. 1J**). Overall, this pattern is consistent with the presence of some cooperativity in the amplification of external perturbations within the ensemble (*39*). In the second regime with dominant E-I coupling (*k* = 4), however, we observed a different trend. The strength of induced ensembles grew sublinearly with *N*_*p*_, plateauing when around ∼30% of all excitatory neurons were perturbed, and dropping for larger fractions of stimulated neurons (**Fig. 1M**). These results indicate that, contrary to the prima facie assumption, increasing the number of targeted neurons by external perturbations may not always lead to formation of stronger neuronal assemblies in E-I networks.

We observed similar results for networks with higher ratio of excitatory to inhibitory neurons (**Fig. S1A**), in networks with sparse connectivity of E-E connections (**Fig. S1B**), and when the network connectivity was random rather than specific (**Fig. S1C**). Our results also hold for different variants of Hebbian rule (**Fig. S2**). We focused our analysis in the following sections on randomly connected networks with covariance-based learning rule (see **Methods**).

### Transition from cooperative to suppressive regimes

To gain further insights into the formation of assemblies in different regimes, we analyzed how the average strength of individual synapses change as a function of parameters of perturbation (**Fig. 2A-F**). Within the ensemble of targeted neurons, we plotted the average potentiation of synapses for the two regimes (**Fig. 2A,D**). For networks with weak E-I coupling, increasing N and T both enhanced the average induction (**Fig. 2A**), indicating that the recurrent interactions amplify the strengthening of ensembles. Such enhancement of potentiation per synapse combined with the increase in the number of presynaptic sources leads to the supralinear potentiation of the weights of the assemblies, as we observed before (**Fig. 1J**). Networks with dominant E-I coupling, on the other hand, showed a suppressive behavior per synapse: average enhancement of synapses was *decreased* for larger perturbation sizes (**Fig. 2D**). Combination of this suppressive effect with the increase in the number of presynaptic sources led to a sublinear growth of the total potentiation of the ensemble, as we observed before (**Fig. 1M**). Similar dependence on the induction size was also observed for more biologically realistic implementation of networks with spiking neurons in different E/I regimes (**Fig. S3**).

**Fig. 2.**
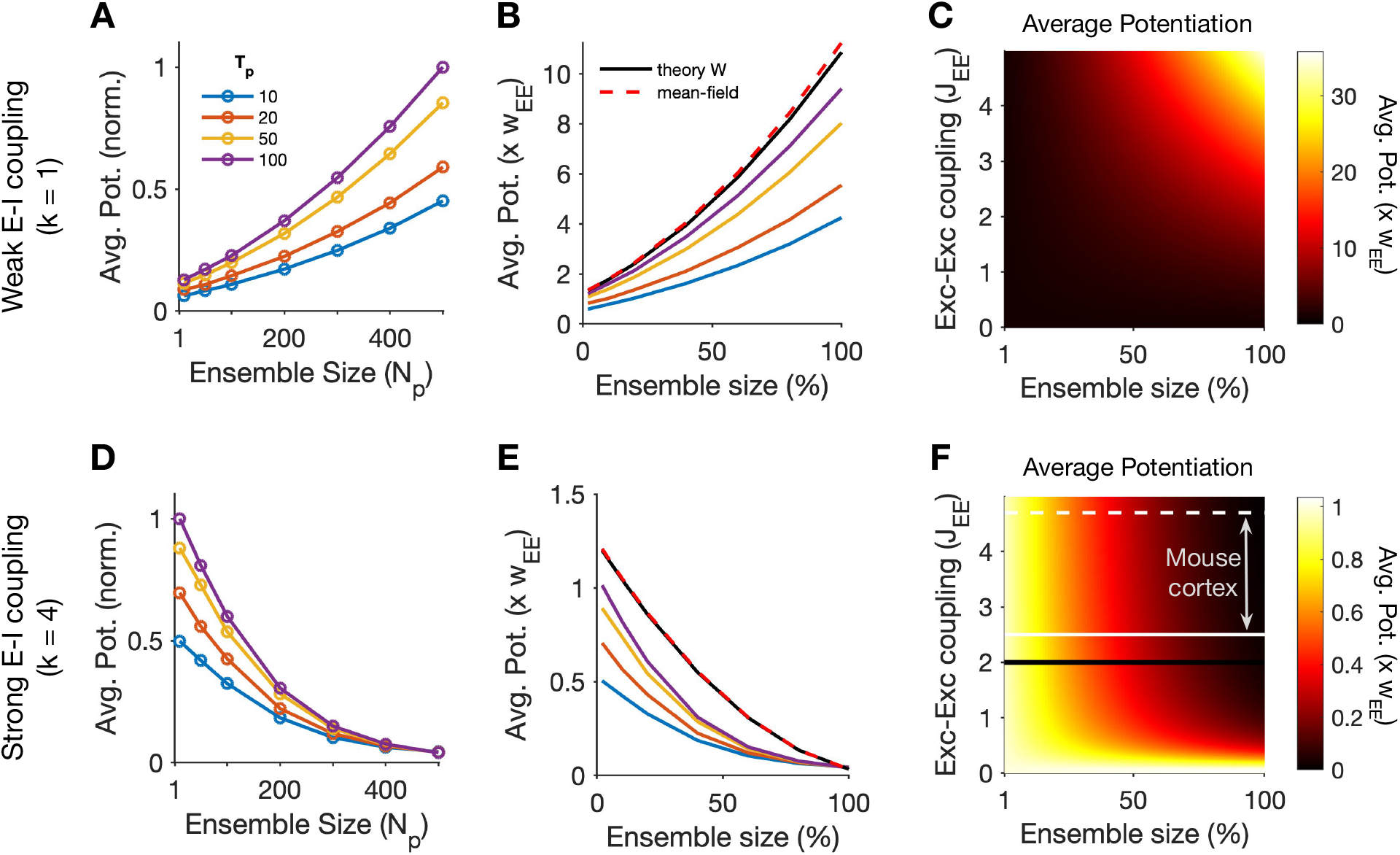
Transition from cooperative to suppressive regimes. (**A**) Average potentiation (Avg. Pot.) of individual synapses within the assembly of perturbed neurons (cf. **Fig. 1J,M**) for different ensemble sizes (*N*_*p*_) and temporal profiles of perturbation (*T*_*p*_), normalized to the maximum. (**B**) The values of average potentiation relative to the average E-E weights in the network (*w*_*EE*_), compared with the theoretical values obtained from linearized dynamics of the network based on its weight matrix (theory W) and from the mean-field analysis (dashed line) (see **Methods** for details). The results of simulations for larger *T*_*p*_ values converge to the theoretical values inferred from *W*, which in turn match with the mean-field analysis. Ensemble size is expressed as a fraction of total E neurons in the network (*N*_*p*_/*N*_E_, where *N*_E_ = 500). Other parameters the same as **Fig. 1**. Networks are in the weak E-I coupling regime (*k* = 1). (**C**) Average potentiation relative to *w*_*EE*_ calculated from the mean-field analysis for different combination of network E-E coupling (*J*_*EE*_ = *N*_E_ *w*_*EE*_) and the size of neuronal ensembles as a fraction of the total size of the network (*N*_*p*_/*N*_E_). (**D-F**) Same as (A-C) for induced ensembles in networks with strong E-I coupling (*k* = 4). The black line in (F) corresponds to previous simulations in (D,E) with *J*_*EE*_ = 2. White lines indicate the range of *J*_*EE*_ estimated in mouse cortical networks, with the solid and dashed lines corresponding to the mode (*J*_*EE*_ = 2.5) and the median (*J*_*EE*_ = 4.7) of the estimated values (*24*).

Distinct dependence of the induction on the size of perturbed neurons was predicted by our theoretical analysis (**Fig. 2B,E**), which calculated the potentiation of synapses from the covariance of response changes resulting from the dynamics of the network, given the initial weight matrix (see **Methods**). The results of network simulations approached the theoretical limit for larger values of *T*_*p*_, reflecting the fact that our analysis considers the stationary state responses of the networks and ignores the temporal dynamics of the transients, which become more dominant in perturbations with smaller *T*_*p*_. Consistent with this reasoning, inferring the potentiation from very short transient responses almost abolished the dependence on the perturbation size; conversely, very large values of *T*_*p*_ matched well with the theoretical prediction and numerical simulations (**Fig. S4**). These results suggest that perturbation protocols employing very fast alternating pulses may fail to reveal the effect of network dynamics on plasticity, as recurrent E/I interactions may not emerge at such short time scales.

To further understand the behavior of networks in different regimes, we developed a mean-field analysis based on the average behavior of the perturbed and nonperturbed subpopulations (see **Methods**). The result of the mean-field analysis matched well with the previous theoretical analysis inferred from the detailed weight matrix of the network (**Fig. 2B,E**). Employing the mean-field analysis, we could scan a large parameter space of arbitrarily large-scale networks with different E-E coupling and different fractions of targeted neurons (**Fig. 2C,F**). The results suggested that for weak E-I coupling, increasing both parameters increases the average potentiation of synapses (**Fig. 2C**), consistent with the conventional assumption that stronger excitatory connections and larger induction sizes both enhance the potentiation of assemblies. For strong E-I coupling regimes, on the other hand, we observed a different behavior. Apart from a small part of the parameter space for very weak E-E coupling, the opposite dependence on *N*_*p*_ was observed, namely suppression of the average potentiation for larger perturbation sizes (**Fig. 2F**). This relationship became steeper for stronger E-E couplings and particularly held for the range of E-E couplings recently estimated in the mouse cortical networks (*24*). Such networks are thus predicted to show unintuitive dependence on the size of perturbation, if they are operating in strong E-I coupling regimes, as suggested for the functional connectivity of the mouse primary visual cortex (*19, 34*).

### Strength versus specificity of induced ensembles

Our results so far indicated that cooperativity in the formation of neuronal ensembles emerges in networks with weaker E-I couplings, and that this changes to suppressive behavior in networks with stronger E-I interactions. Neuronal ensembles are thus expected to emerge faster and stronger in the former regime compared to the latter. But how specific would the outcome of the induction be in each regime? To answer this, we quantified the selectivity of assembly formation by comparing the strength of presynaptic weights of perturbed neurons arising from neurons within and outside the assembly (**Fig. 3A**). If the outcome of induction is specific, the potentiation of weights remains confined to connections within the intended assembly. On the other hand, perturbation of the targeted neurons can lead to off-target effects, causing a potentiation of synapses from (or to) outside the assembly, thus creating a nonspecific potentiation.

**Fig. 3.**
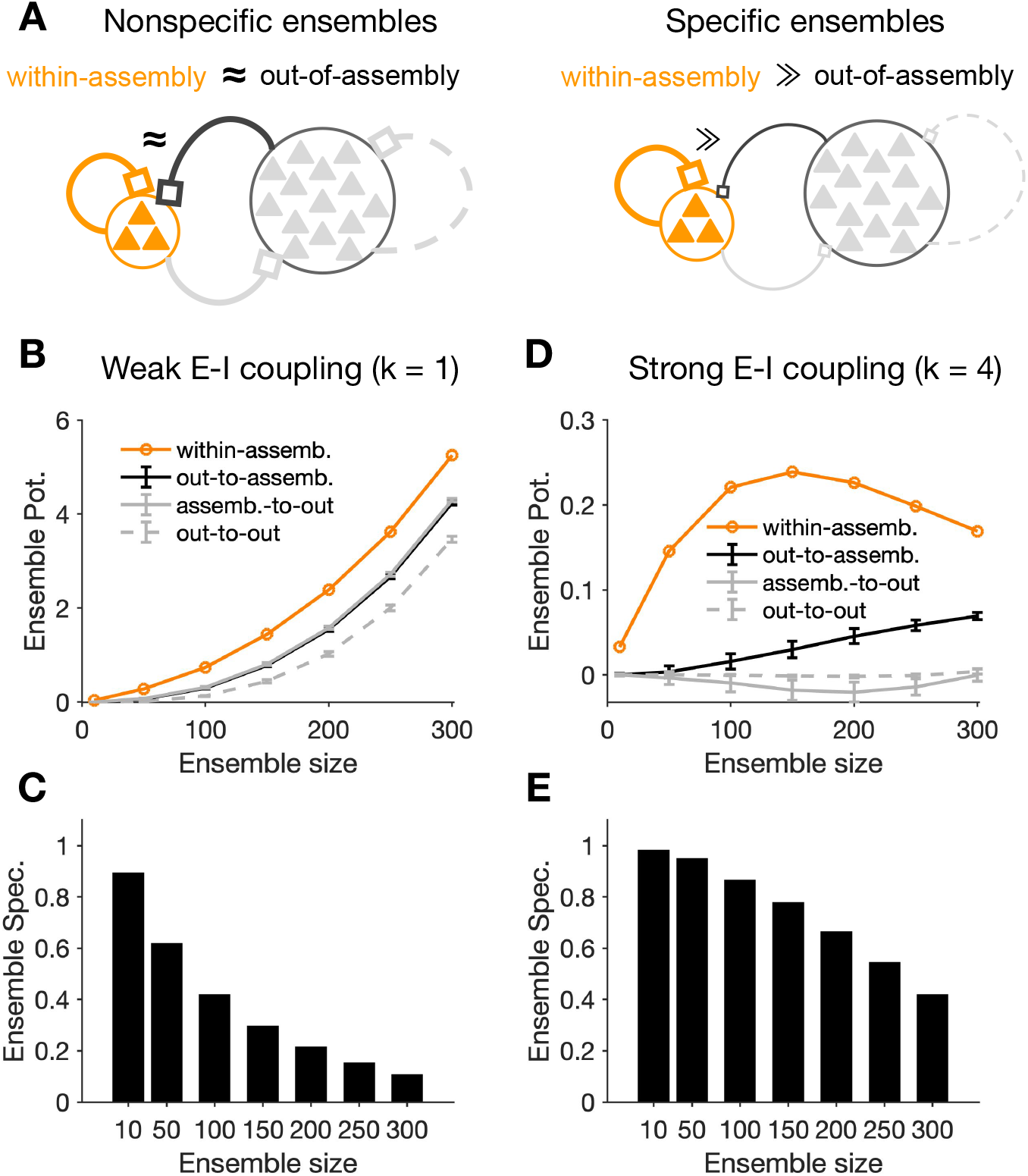
Specificity of assembly formation in different regimes of E/I balance. (**A**) The outcome of induction can be nonspecific (left), if the within-assembly potentiation of weights is accompanied by a substantial potentiation of connections originating from outside the assembly, or specific (right), when the potentiation of weights remains predominantly within the intended ensemble. (**B**) Potentiation of presynaptic connections within the assembly (orange) versus those from the assembly to outside (assemb.-to-out; gray), from outside to the assembly (out-to-assemb.; black), and within the outside neurons (out-to-out; gray dashed), respectively. *T*_*p*_ = 50 and inductions is in the weak E-I coupling regime (*k* = 1). Ensemble potentiation is calculated as the average (across postsynaptic neurons) of the sum of connection weights from all presynaptic sources (cf. **Fig. 1J,M**). For each *N*_*p*_, out-of-assembly potentiation is calculated for 100 randomly selected pools of neurons other than, but with the same size (*N*_*p*_) as, the perturbed neurons. Line and error bars show the average and std across the pools, respectively. (**C**) Ensemble specificity (Spec.) quantifies the specificity of induced ensembles for different sizes of perturbed neurons. It is calculated as (*E*_*w*_ – *E*_0_)/(*E*_*w*_ + *E*_0_), where *E*_*w*_ and *E*_0_ are the average within and out-of-assembly (assemb.-to-out) ensemble potentiation in (B), respectively. Ensemble specificity drops for larger ensemble sizes, reflecting the fact that within-assembly potentiation of weights is accompanied by a substantial potentiation of connections from outside. (**D,E**) Same as (B,C) for neuronal ensembles forming in networks with strong E-I coupling (*k* = 4). Out-of-assembly potentiation grows much slower than within assembly potentiation initially until the latter plateaus and starts to drop (D), leading to a high ensemble specificity for initial ensemble sizes (E).

For networks with weak E-I coupling, strong within-assembly potentiation was accompanied by a substantial out-of-assembly potentiation of weights, resulting in a significant drop in ensemble specificity for large perturbation sizes (**Fig. 3B,C**). However, potentiation of connections from outside the ensemble grew much slower for networks with strong E-I coupling, leading to an optimal size of induction where the strongest potentiation had a high induction specificity (**Fig. 3D,E**). We observed qualitatively similar results for different variants of the Hebbian plasticity rule (**Fig. S5**). These results show that stronger potentiation of assemblies in networks with weak E-I coupling comes at the price of losing ensemble specificity, as the relative potentiation of within-assembly to out-of-assembly weights decreases for larger perturbation sizes. Strong E-I interaction hampers the potentiation of ensembles, but leads to a more selective formation of neuronal assemblies, which is more robust to the size of perturbations.

In randomly connected networks, the distribution of weight changes from non-assembly neurons is random, irrespective of different mean values in different regimes (**Fig. 3B,D**). However, preexisting wiring in the network may guide the process of induction (*40*) and lead to a non-random distribution. Connectivity between neurons are in fact reported to be organized according to their functional properties (*22, 23, 41, 42*). We therefore asked how the modulation of out-of-assembly connections depends on the initial network structure in networks with some non-random (specific) connectivity structure. Specific connectivity was implemented by modulating the connectivity weights in the network to have stronger connections between pairs of neurons with similar functional properties, which was assumed to be a one-dimensional feature (e.g. preferred orientation) here (cf. **Fig. 1I,L** and **Methods**).

In weak E/I regime (*k* = 1), we found feature-specific potentiation, namely out-of-assembly connections potentiated more for neurons with similar functional features as the perturbed neurons (**Fig. S6A**). This result suggests that preexisting connectivity in this regime interferes with the potentiation of the induced ensemble. For networks with strong E/I interactions (*k* = 4), on the other hand, we observed an opposite trend: neurons closer in the functional space experienced, on average, a larger depression of their weights to the ensemble (**Fig. S6B**). Such feature-specific depression of weights can increase the specificity of induction, by suppressing the strong presynaptic connections that are irrelevant to the intended ensemble. Different regimes of E/I can therefore support different modes of assembly formation with regard to preexisting structure of the network, with weak E-I coupling regimes promoting the influence of the previous connectivity, and strong E-I coupling enabling a more efficient “rewriting”.

Both feature-specific potentiation and depression were absent when initially perturbed neurons were chosen randomly, independent of their preferred orientations (**Fig. S6A,B**). These results therefore argue that different regimes of recurrent interaction as well as different patterns of induction can lead to distinct outcomes of plasticity. Note that, while we assumed similar properties for all neurons, accounting for more biologically realistic receptive fields of E/I neurons (*34*) and their connections (e.g. broader selectivity and connectivity of inhibition (*43–46*)) could lead to a center-surround pattern of out-of-assembly plasticity, with potentiation and depression for highly similar and less similar connections, respectively (via-a-vis center-surround patterns of influence resulting from neuronal perturbations in the visual cortex (*19, 34*)).

Neuronal ensembles are suggested to be involved in subnetwork-specific recovery of responses following input deprivation (*47*). We therefore asked how this process can be guided by specificity of resulting assemblies in different regimes. We reduced the feedforward input to a fraction of neurons in the network (comprising distinct subnetworks, A and B), and studied how correlated external activation of a subset of them (subnetwork A) can lead to recovery. In both regimes, neurons in subnetwork A potentiated their recurrent weights, which can counteract the lack of feedforward drive after input deprivation (**Fig. S6C,D**). While this potentiation happened exclusively within subnetwork A in networks with strong E-I coupling (**Fig. S6D**), recovery in weak E-I coupling regimes was also accompanied by potentiation of connections from other E neurons (**Fig. S6C**). Specifically, the reciprocal connectivity between subnetworks A and B was potentiated in the weak E-I regime, while it was depressed in the strong E-I regime (**Fig. S6C,D**). These results therefore suggest that strong E-I interactions can shape the specificity of formation of neuronal assemblies in the network and their subsequent recovery following input deprivation.

### Speed and specificity of assembly formation

In the previous analyses, we focused on how response changes resulting from perturbations in different dynamic regimes guide network-wide plasticity and formation of assemblies (**Fig. 1C**). Such unidirectional effects of dynamics on plasticity might in fact be pertinent to initial stages; in later stages, however, weight changes would in turn shape the network dynamics (although with a slower time course). To fully analyze the dynamic evolution and growth of neuronal assemblies in cortical networks, we therefore need to consider this closed-loop interaction of dynamics and plasticity (**Fig. 4A**).

**Fig. 4.**
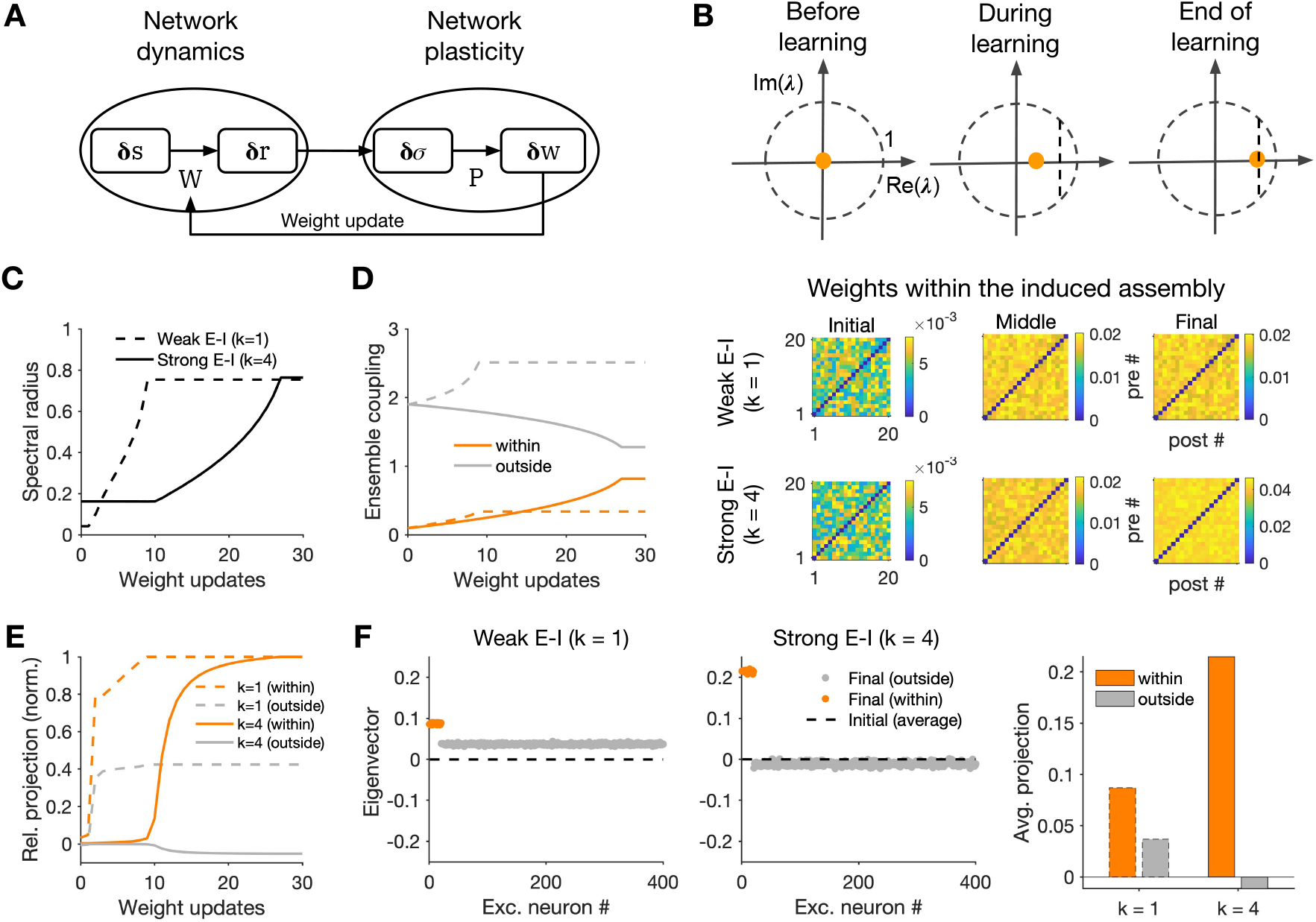
Growth of ensembles in networks with recurrent interaction of dynamics and plasticity. (**A**) Closed-loop interaction of network dynamics and network plasticity underlying the formation and growth of neuronal ensembles. Network dynamics governed by the weight matrix (W) determines the input-output responses to external perturbations, which in turn shape the structure of covariances in the network. Network plasticity (P) guided by the resulting covariance patterns determines the weight changes and updates, on a slower time scale, the weight matrix, which, in turn, modifies the network dynamics. (**B**) Upper: The spectral radius of the network, denoting the growth of the maximum eigenvalue of the weight matrix (λ_0_), at different steps of weight update. To avoid instability of the network dynamics (*λ*_0_ > 1), the learning is stopped before *λ*_0_ reaches a threshold close to 1 (vertical dashed line). Lower: Sample weight matrices of the perturbed neurons at different stages of weight updates, for networks in different E/I regimes. *N*_*p*_ = 20, *T*_*p*_ = 50, other parameters the same as networks in **Fig. 2**. (**C**) Evolution of the spectral radius in different regimes. Neuronal ensembles in networks with weak E-I coupling (*k* = 1) reach the upper bound on the spectral radius faster, and their growth is therefore limited sooner, than network with *k* = 4 (cf. (B)). (**D**) Ensemble coupling within (orange) and from outside (gray) the ensembles (cf. **Fig. 3B,D**) at different steps of weight update (dashed: *k* = 1, solid: *k* = 4). (**E**) Relative projection of the eigenvector (*v*_0_) corresponding to the largest eigenvalue (*λ*_0_) of the network over neurons within (orange) and outside (gray) the ensemble for networks with *k* = 1 (dashed) and *k* = 4 (solid). It is calculated as the average real part of the entries corresponding to perturbed and non-perturbed neurons, respectively, and normalized by the maximum value for each regime. (**F**) Left: Distribution of the real part of the largest eigenvector (*v*_0_) over excitatory neurons at the end of learning. For comparison, the dashed line shows the average value (across all excitatory neurons) of the initial, random distribution before induction. Right: Average projection of the final eigenvector (shown on the left) over excitatory neurons within and outside the assembly.

To study this, we repeated our previous perturbation protocols while updating the weight matrix of the network in incremental steps. The weight matrix was updated in time intervals of Δ*T*_*w*_, while between the updates the weight matrix (*W*) was kept constant and determined the network dynamics. Note that Δ*T*_*w*_ is much larger than the time scale of network integration (τ), which is justified by the separation of time scales of dynamics and plasticity (*48*). We used a Hebbian rate-based covariance rule to update the weights (see **Methods**). To ensure the stability of network dynamics, the weight update is performed at each stage only if the largest eigenvalue of the weight matrix (or its spectral radius) does not grow more than a value close to, but smaller than, one (**Fig. 4B**; **Methods**). Different mechanisms can be employed to ensure such stability, e.g. hard bounds for the weights, weight normalization, synaptic scaling or inhibitory stabilization (*38, 49, 50*), but our analysis here remains agnostic about the nature of this mechanism.

Growth of the spectral radius provides a proxy for the speed of learning in different regimes (**Fig. 4C**). The spectral radius grew much faster for networks with weak E-I coupling (*k* = 1), indicating a faster strengthening of weights in this regime. Evolution of the spectral radius was similar to the fast strengthening of weights within the induced neuronal ensemble in this regime (**Fig. 4D**). The fast assembly growth was, however, accompanied with the fast potentiation of out-of-assembly connections (**Fig. 4D**). The evolution of neuronal assemblies in the strong E-I coupling regime (*k* = 4), on the other hand, was slow and specific: both the spectral radius and within-assembly weights grew much slower, but this was accompanied by weakening of connections from outside the ensemble, leading to specificity of assembly formation (**Fig. 4C,D**). Different patterns of growth of neuronal assemblies in different regimes can be explained in terms of the eigenvector corresponding to the largest eigenvalue of the network at each stage (**Fig. 4E-F**). If the connections within the induced ensemble are mainly potentiated over time, the eigenvector will have specific projections over the perturbed neurons. Nonspecific growth would, on the other hand, translate to nonspecific projection of this eigenvector over perturbed and unperturbed neurons. In fact, networks with weak and strong E-I couplings show, respectively, such nonspecific and specific projections during learning (**Fig. 4E**) and at the end of it (**Fig. 4F**). Thus, although the largest eigenvalue of the network grows faster in weak E-I regimes (**Fig. 4C**), its corresponding eigenvector does not remain confined to perturbed neurons (**Fig. 4E**), indicating that within-assembly potentiation of weights is accompanied by potentiation of connections from outside the ensemble (**Fig. 4D**). The growth of eigenvalue in strongly coupled E-I regimes is slower, but the corresponding eigenvector and the potentiation of weights remain specific to perturbed neurons, ensuring a specific formation of neuronal assemblies (**Fig. 4C-F**).

### Pattern completion and different regimes of recall

We also studied how neuronal ensembles emerging in each regime show pattern completion (**Fig. 5**). At the end of learning (**Fig. 4B**), we partially activated the neurons within the induced ensemble and measured the response of other neurons, which were not directly activated by the external stimulation (**Fig. 5A**). Neuronal ensembles formed in both regimes showed pattern completion when half of their neurons were activated (**Fig. 5B,D**). We further quantified the strength of pattern completion for different fractions of partial activation. This was calculated by comparing the average response of the nonactivated and activated neurons within the ensemble (see **Methods**). Both networks showed comparable pattern completion curves within the ensemble, with even small fractions of activation eliciting significant responses in nonactivated neurons (**Fig. 5C,E**).

**Fig. 5.**
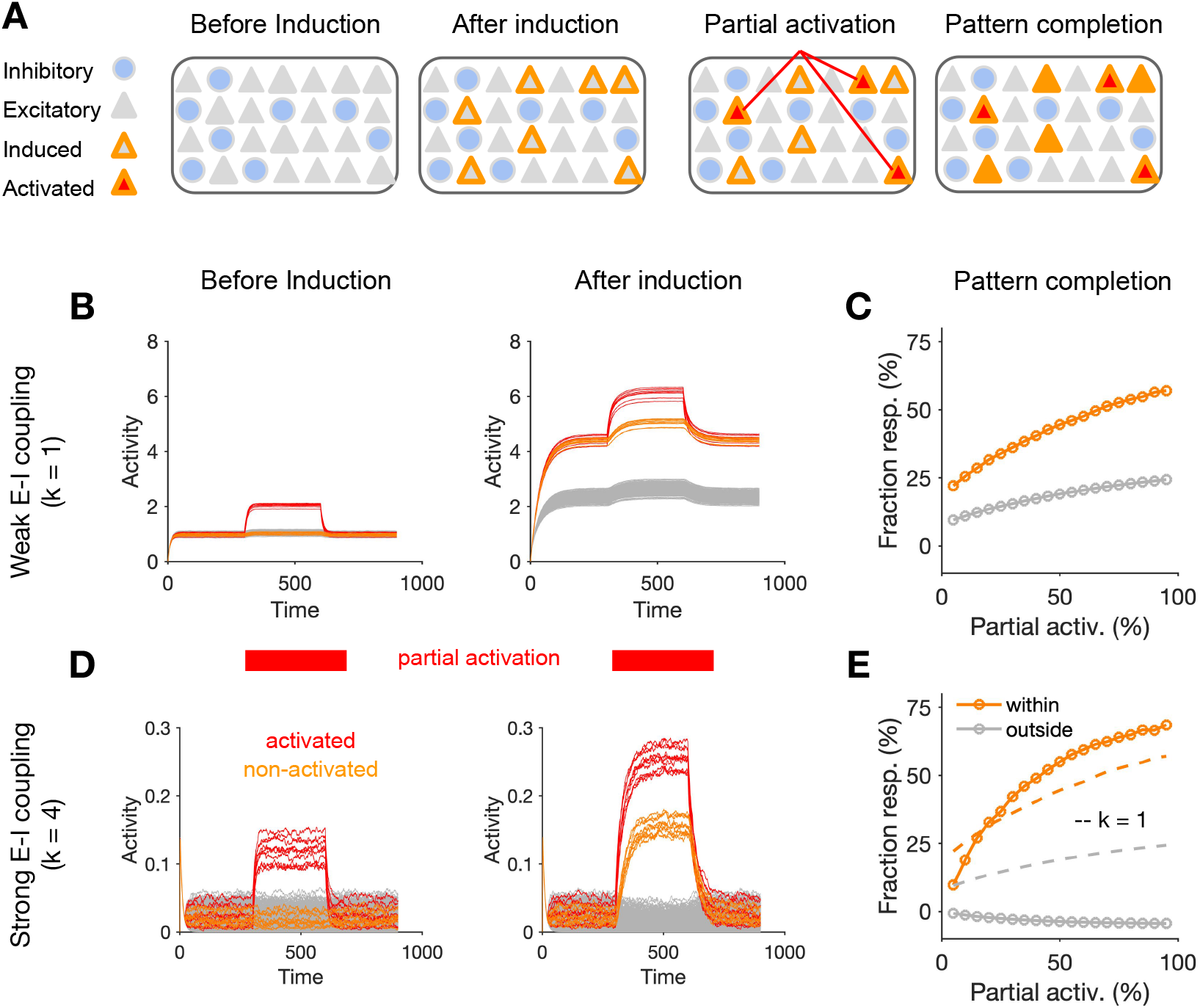
Pattern completion in neuronal ensembles emerging in different E/I regimes. (**A**) Pattern completion is triggered by partial activation of neurons within an induced ensemble and evaluating the response of other, nonactivated neurons. (**B**) Response of the network with weak E-I coupling (*k*= 1) in the baseline with and without partial activation of the assembly. Left: before formation of ensemble (before induction); Right: after induction, with updated weights at the end of learning, as described in **Fig. 4**. Half of the neurons in the induced ensemble (10 out of 20, shown in red) are stimulated by extra perturbations, and the effect on other excitatory neurons within (orange) and outside (gray) the ensemble is evaluated. (**C**) Pattern completion curve describing the degree of pattern completion (quantified by Fraction resp.) as a result of partial activation of the ensemble (Partial activ.). Fraction resp. is calculated as the average response change of the nonactivated neurons (within (orange) and outside (gray) the assembly, respectively) divided by the average response change of activated neurons. Response changes are measured relative to the respective baseline activity of each neuron before partial activation. Note that this is a conservative measure for quantifying pattern completion, which only reaches 100% when all non-activated neurons reach the same level of activity as activated ones. (**D,E**) Same as (B,C) for networks with strong E-I coupling (*k*= 4). The pattern completion curve for *k*= 1 is copied in (E) for comparison (dashed lines). Note that the strength of external perturbations is larger for *k*= 1, to adjust for the higher baseline activity of the network (see **Methods** for details).

However, pattern completion was more specific in networks with stronger E-I couplings. In the network with strong E-I coupling (*k* = 4), recurrent activation of nonactivated neurons remained specific to the induced ensemble (**Fig. 5E**). In contrast, in the network with weaker E-I coupling (*k* = 1), recurrent interactions also elevated the activity of neurons outside the assembly, leading to some degree of nonspecificity in pattern completion (**Fig. 5B,C**). Taken together, these results suggest that neuronal ensembles forming in networks with weak or strong E-I coupling may enable computations with different speed-accuracy trade-off. If such neuronal ensembles guide behavior, they may, in turn, lead to different regimes of cognitive processing.

### Behavioral performance associated with neuronal assemblies in different E/I regimes

We next studied how neuronal assemblies can contribute to behavioral performance in different regimes of recall (**Fig. 6**). We simulated the development of two neuronal assemblies (A and B), associated with two stimuli corresponding to distinct behavioral contexts (**Fig. 6A**, left). The association was established in induction sessions, where neurons belonging to each assembly were perturbed (similar to protocols described in **Fig. 4**; *N*_*p*_ = 20). The behavioral performance was assessed in recall sessions (**Fig. 6B**), where a fraction of neurons (5/20) in each assembly (A or B, respectively) was stimulated. The capacity of the network to detect the presence of a context was assayed by quantifying the “recall strength” of the respective assembly (see **Methods**). The performance of the network to distinguish between the two behavioral contexts was quantified by calculating a “discriminability index” (*d′*), which compared the response of a given assembly to its target and distractor contexts (see **Methods** for details).

**Fig. 6.**
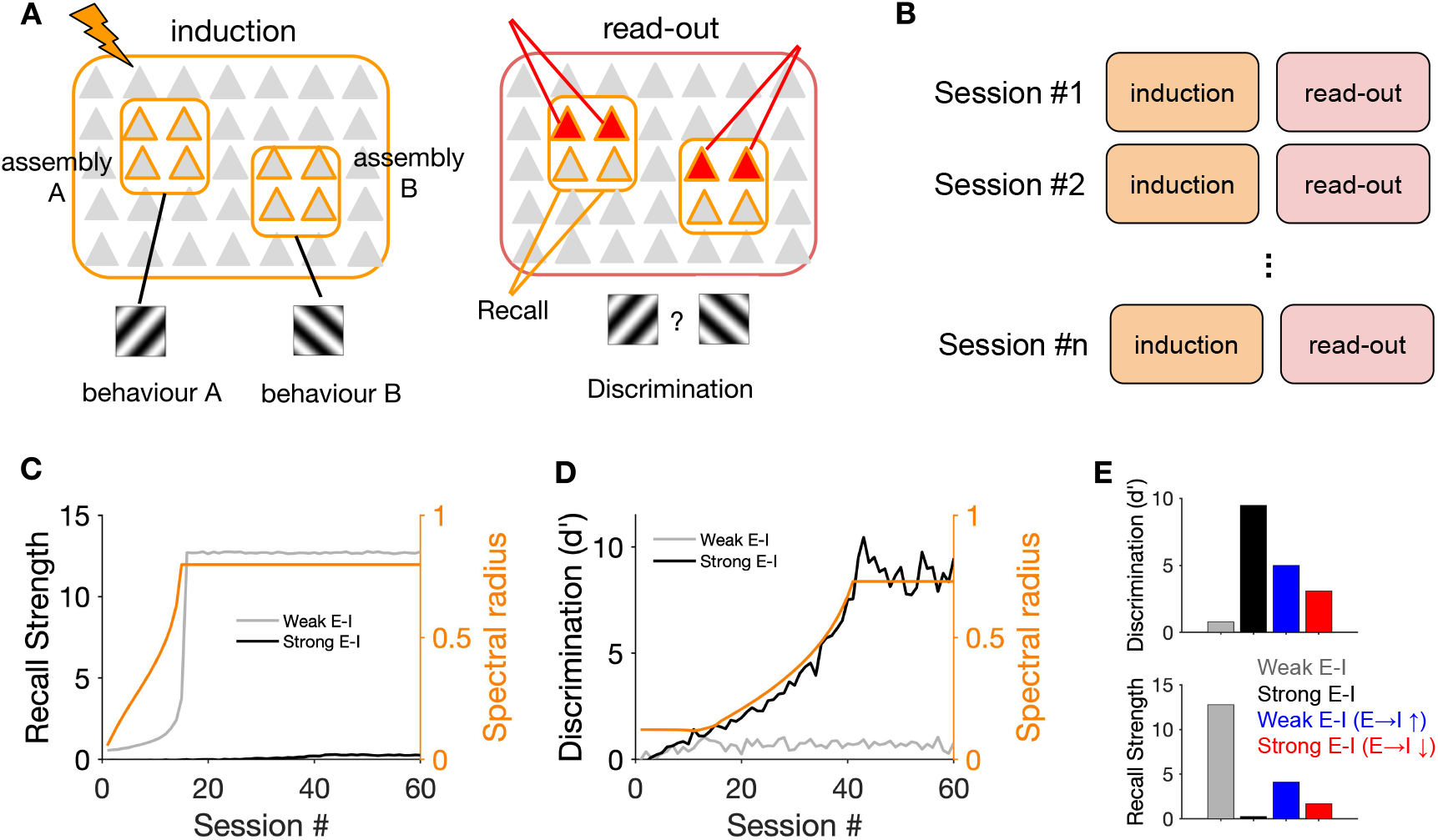
Performance of neuronal assemblies in behavioral tasks in different regimes of recall. (**A**) Left: In networks with different regimes of E-I coupling, two neuronal assemblies (A and B) are induced to represent two behavioral contexts (A and B, respectively). Right: Performance of the network in behaviorally relevant tasks are evaluated from the read-outs of ensemble responses to partial triggers. (**B**) The read-outs are evaluated after each induction session, and the weights are updated in the next session following the same procedure described in **Fig. 4**. *N*_*E*_ = *N*_*I*_ = 200. (**C**) Recall strength is quantified in each session to evaluate the capacity of the ensemble (*N*_*p*_ = 20) to detect its respective context. It is calculated as the average increase in the activity of non-triggered neurons (15/20) in the assembly, when a small fraction of neurons (5/20) are stimulated to trigger the corresponding behavior. The evolution of the spectral radius in the weak E-I regime (*k*= 1) is plotted on the right y-axis for comparison. (**D**) A discriminability index is calculated at each read-out session to evaluate the capacity of the ensembles to distinguish between the two contexts (see **Methods**). The evolution of the spectral radius in the strong E-I regime (*k*= 4) is plotted on the right y-axis for comparison. (**E**) Discrimination and recall strength for networks with weak E-I (gray) and strong E-I (black) coupling at the end of learning. The results are compared with networks with weak E-I coupling when the recruitment of inhibition is boosted via E→I connections (inhibitory modulation; blue), and to networks with strong E-I coupling where E→I connections are weakened (disinhibitory modulation; red). These modulations enhance discriminability in networks with weak E-I, and increase the recall strength in networks with strong E-I coupling regimes, respectively.

Networks with weak E-I coupling showed a very swift increase in recall strength, which matched with the quick growth of their spectral radius (**Fig. 6C**). This shows that neuronal assemblies in this regime can amplify a weak stimulation of a small fraction of their neurons, providing a substrate for fast and strong recalls. In comparison, recall strength was much weaker and rose up much more slowly in networks with strong E-I coupling (**Fig. 6C**). Neuronal ensembles in the latter regime had, however, a significant advantage in discriminating between the two contexts (**Fig. 6D**). While the initial enhancement of discriminability (*d′*) plateaued in weak E-I regimes, neuronal assemblies in strong E-I regimes improved their discrimination capacity for much longer and to much higher values, matching the slower growth of their spectral radius (**Fig. 6D**).

These results suggest that neuronal ensembles emerge slower in inhibition-dominated regimes and enable fine downstream readout discriminations, while the assemblies forming in weaker E-I regimes can be suited for faster but less specific cognitive tasks. Modulating E-I balance in the network, for instance by top-down mechanisms (e.g. via vasoactive intestinal peptide (VIP)-positive neurons) (*51–53*) can, therefore, provide a powerful tool to control different modes of learning (*54*). We tested this in our networks and found that modulating E→I coupling bidirectionally modulated learning: increasing E→I coupling in weak E-I networks increased discrimination (and decreased the recall strength), while decreasing E→I coupling in strong E-I networks increased recall strength (and reduced discrimination) (**Fig. 6E**). Different modes of learning and induction can therefore be achieved by general modulation of the network.

### Dynamic transition between different regimes resulting from E-I plasticity

In our networks so far, we only allowed E-E synapses to be plastic, and studied the effect of E-I interactions on this plasticity by changing static E-I weights in different E/I regimes. Such regimes may not be static, however, and can be dynamically modulated, as we discussed in the previous section. In addition to external mechanism like top-down modulation, plasticity of E-I connections within the network can also intrinsically change the E/I regime (*55, 56*). Different plasticity rules of subtypes of inhibitory neurons indeed shape dynamics and learning in different manners (*57–59*). We therefore studied how E-I plasticity may contribute to different regimes of induction by extending our model networks and allowing E→I and I→E synapses to be governed by Hebbian rules based on covariance of responses, similar to E-E weights (cf. **Fig. 4**).

Specifically, we asked if combining E-E and E-I plasticity can enable a network with weak E-I coupling to dynamically transition to a strong E-I regime of induction (**Fig. 7A**). Induction of neuronal assemblies in such a network indeed led to a strong potentiation of perturbed excitatory neurons (**Fig. 7B**, left); this potentiation was much weaker when E-I plasticity was inactive (**Fig. 7B**, right and **Fig. 7D**). Potentiation of E-E ensembles was anticorrelated with the average activity of the networks: networks with E-E and E-I plasticity in fact decreased their baseline activity over the course of learning, while the baseline activity increased for networks in which E-I plasticity was inactive (**Fig. 7C**). Sparsification of activity was a result of potentiation of E-I coupling, which put the network in a more inhibition-dominated regime.

**Fig. 7.**
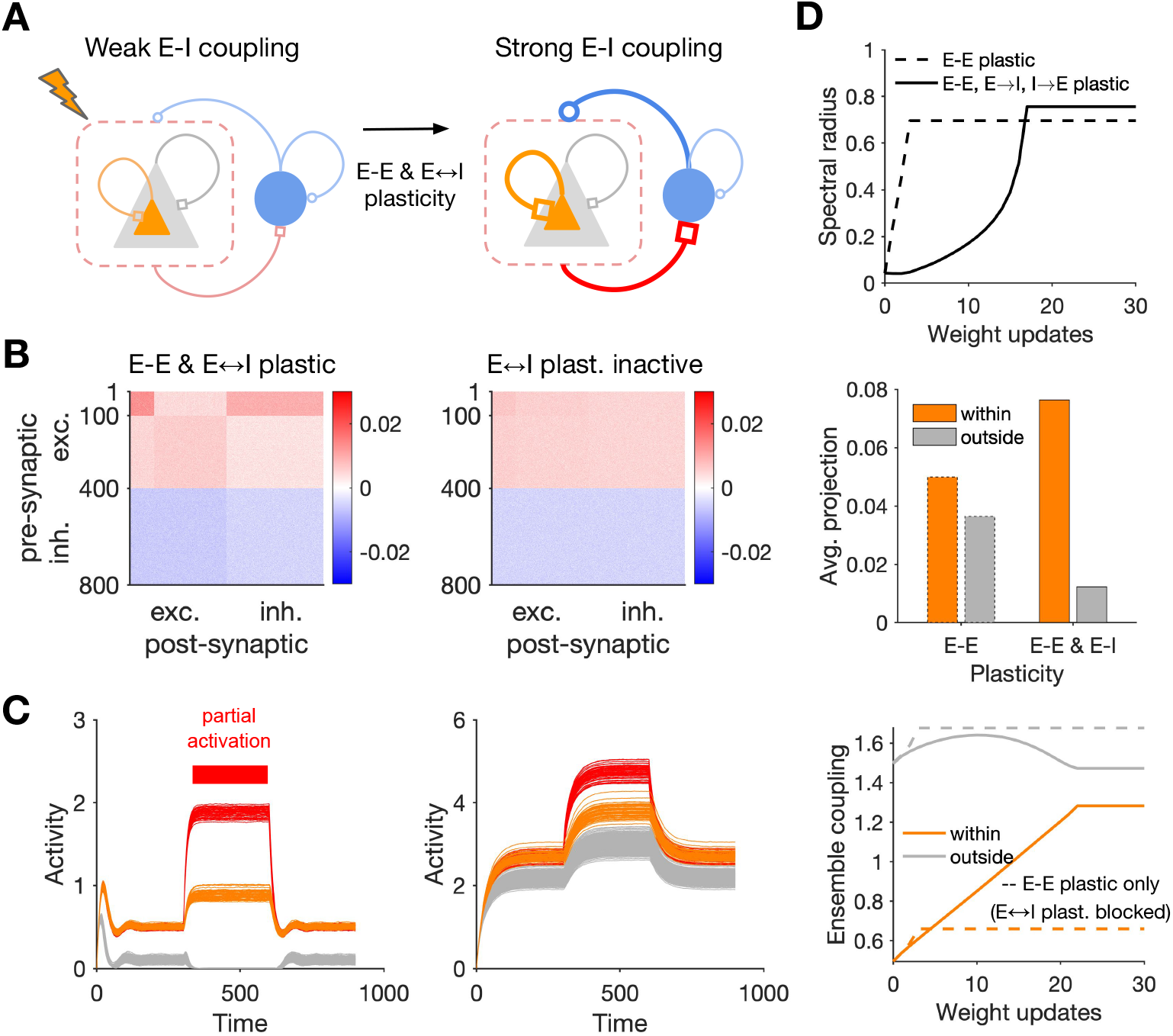
Dynamic transitions between different regimes of assembly formation. (**A**) Schematic of a network with E-E and E-I plasticity before and after induction of assemblies. (**B**) Final weight matrix of the network at the end of learning (similar to the procedure described in **Fig. 4**), in the network where both E-E and E-I weights are plastic (left), compared with the condition where E-I plasticity is blocked and only E-E plasticity remains (right). *N*_*E*_ = *N*_*I*_ = 400, *N*_*p*_ = 100 (perturbed neurons #1-100). (**C**) Pattern completion in networks with E-E and E-I plasticity (left) and only E-E plasticity (right), at the end of learning. (**D**) Growth of the spectral radius (top; cf. **Fig. 4c**), average projection of the largest eigenvector over excitatory neurons (middle; cf. **Fig. 4F**), and evolution of ensemble coupling (bottom; cf. **Fig. 4D**), in the networks with E-E and E-I plasticity (solid lines) and when E-I plasticity is blocked (dashed lines).

The network with E-E and E-I plasticity showed selective pattern completion upon partial activation of neurons in the ensemble (**Fig. 7C**, left), and this selective pattern completion was abolished after inactivating E-I plasticity (**Fig. 7C**, right) (cf. **Fig. 5B,D**). Slow but selective growth of the eigenvector associated with the largest eigenvalue shed light on the slow and selective potentiation of within-ensemble weights, which was in contrast to fast and nonspecific formation of assemblies when E-I plasticity was blocked (**Fig. 7D**; cf. **Fig. 4C-F** for different E-I regimes). Our results also hold for another implementation of E-I plasticity based on pre- and post-synaptic covariances (**Fig. S7**). Taken together, these results suggest that E-I plasticity can enable the network to dynamically transition between different regimes of induction and learning.

## Discussion

We studied how different patterns of perturbations can induce neuronal assemblies in large-scale balanced networks. Our results revealed different regimes of induction for the spectrum of excitation-inhibition balance. In particular, we found that increasing the size of perturbed neurons may not always lead to more potentiation. Induced assemblies in regimes with dominant E-I coupling exert a potent lateral inhibition, which suppresses the activity of neurons and the potentiation of their respective connections. This would also apply to connections within the intended assembly in a recurrent manner, leading to a sublinear growth of the total weights. Although hampering the strength of plasticity, the mutual inhibition increases the specificity of ensembles, by suppressing the nonspecific potentiation of connections.

Our results therefore suggest that inhibition can gate and modulate the specificity of induction. It slows down the formation and growth of neuronal assemblies, but ensures that perturbation-induced learning remains specific to intended ensembles, and does not lead to off-target effects. The selectivity remained the same for various sizes of induced ensembles, suggesting that regimes with dominant E-I coupling also guarantee the size-invariance of plasticity. We could further show, in our model behavioral experiments, that such inhibition-dominated regimes are best suited for fine discrimination tasks, which rely on selectivity of neuronal assemblies. It would be interesting to test these predictions in future experiments, by modulating E/I balance (*60*) in cortical networks and measuring the discriminability of behavioral responses.

When excitation was predominant, induction was fast and strong in our network models, but did not remain specific to the induced assembly. Due to an indirect recurrent recruitment of other neurons in the network, connections from outside the assembly also strengthened. This compromised the specificity of pattern completion by neuronal ensembles, and it reduced the capacity of the network to discriminate between behavioral contexts represented by different ensembles. This regime might, instead, be better suited for crude detection tasks, and can provide a substrate for generalization to other assemblies and beyond a specific context.

Which regimes of induction are more pertinent to the regimes in which cortical networks operate? Functionally, network with strong E-I coupling can provide a natural substrate for recent behavioral findings, which suggest that mutual inhibition of neuronal ensembles can underlie their selective responses (*61, 62*). In terms of connectivity, the key ingredient of the strong E-I coupling regime has been observed in many cortical regions, where a dense and strong connectivity between pyramidal cells and different subtypes of interneurons, including parvalbumin-positive (PV+) and somatostatin-positive (SOM+) cells, has been reported (*63, 64*). In the mouse primary visual cortex (V1), for instance, both pyramidal cell (PC)-to-PV and PV-to-PC connections are an order of magnitude larger than PC-PC connections (*20*). This regime is also consistent, in terms of dynamics, with recent results from single-neuron optogenetic studies (*19*): the prevalence of suppressive effects reported in the experiments only emerges in networks with dominant E-I coupling (*34*). It is therefore likely that mouse V1 operates in an inhibition-dominated regime which favors selective (but weak and slow) induction of assemblies.

This regime might be relevant to other cortices, too. Strong local excitatory-inhibitory coupling has been also reported in other areas, including mouse somatosensory and frontal cortex (*63–66*). In the mouse barrel cortex, optogenetic stimulation of ∼100 excitatory neurons induced a strong *inhibition* of neighboring excitatory neurons, arguing for an inhibition-dominated regime of activity favoring competition and sparsification (*65*). Optogenetic stimulation induced rapid excitation (at ∼5 ms), which was quickly quenched by inhibition (at ∼10 ms) (*65*), consistent with our results on the emergence of suppressive effects for longer pulses (*T*_*p*_) (cf. **Fig. 1M** and **Fig. S4**). A recent study in the mouse premotor cortex found patterned perturbations of a smaller subset of neurons (<10) to induce more excitatory effects in coupled excitatory neurons, although still a significant number of coupled neurons were inhibited (about one third of the excited ones) (*67*). It would be interesting to explore, in future studies, how such differences arising from stimulation protocols and operating regimes can contribute to differential formation of neuronal assemblies in different cortices.

The operating regime of induction can, in turn, be dynamically modulated across different cortices and layers by different factors, including behavioral states (e.g. transition from anesthetized to awake states (*68*) or stationary versus running (*51*)), neuromodulation and attention. For instance, top-down inputs can disinhibit the local circuitry (via VIP→SOM disinhibition) (*51*), VIP neurons can control different stages of learning by differential recruitment of PV neurons (*54*), and the neuromodulatory suppression of PV cells by SOM neurons is crucial for the onset and closure of the critical period of plasticity (*69*). Our results too suggest that modulating E-I coupling -- either via top-down modulation (**Fig. 6**) or intrinsic plasticity of E-I interactions (**Fig. 7**) -- can provide a potent mechanism to control the formation of neuronal assemblies in different regimes of learning. Plasticity of E→I connections has indeed been shown to be crucial for gating of memory and network plasticity (*70*) (cf. **Fig. 7**). It would be interesting to see how such different network states affect the formation of neuronal assemblies and learning, by studying the effect of perturbation protocols in different regimes.

Future theoretical work is also needed to study different regimes of induction in more realistic conditions. First, following the classical notion of neuronal assemblies as recurrent subnetworks, we focused our analysis on the induction of neuronal assemblies in the spontaneous state, where the effect of recurrent interactions are dominant. However, it would be also interesting to see how the plasticity of feedforward synapses in the evoked state interacts with recurrent connections (*71*). Such evoked state will also amplify the effect of stimulus selectivity of neurons, and hence their preexisting connectivity based on that selectivity, which might in turn guide or limit the induction (*40*). Extension of our model networks to allow for feedforward as well as recurrent plasticity can shed light on these more realistic regimes of network responses.

To obtain computational insights into the basic properties of network-wide plasticity and assembly formation, we focused our analysis on simple models of dynamics and plasticity. Inevitably, many biological mechanisms were absent from our models. It would therefore be important to investigate our results in more realistic networks, including those with more complex single-cell mechanisms like dendritic nonlinearity and plateau potentials, and networks equipped with other rules of plasticity (*55, 56, 72*) (e.g. STDP or voltage-based plasticity rules (*73*)) and homeostasis (*50, 74, 75*). For instance, correlations emerging in spiking networks (*76*) especially in excitation-dominant regimes (*77, 78*) may amplify nonspecific potentiation across the network, when STDP rules are employed. It would also be interesting to study how different subtypes of inhibition and their mutual disinhibition (*79*) affect our results, as well as different frequency bands (*80, 81*) and various rules of inhibitory plasticity (*57, 59, 82*) associated with them. Finally, it would be interesting to extend our work to learning sequential chains (*9*) and tasks.

In summary, our work highlights the importance of studying dynamics of neuronal networks and network-wide plasticity in tandem to cast light on the formation of neuronal assemblies. It suggests that unexpected results may emerge when considering the recurrent interactions within networks of excitatory and inhibitory neurons, and that such effects might be missed by focusing on isolated pairs of neurons detached from their network interactions. As behaviorally relevant learning is ultimately happening in ensembles of neurons embedded in large-scale recurrent networks, it is crucial to understand the effect of the background dynamics on the formation of neuronal assemblies and learning. Here, we developed a computational framework to help with this understanding, which can guide the design of future perturbation protocols.

## Materials and Methods

### Network simulations

Rate-based networks were simulated by the following equations for excitatory (E) and inhibitory (I) neurons:

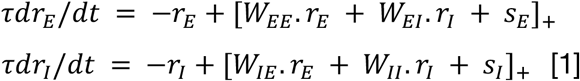

where *r*_*E*_ and *r*_*I*_ are the vectors of firing rates of E and I neurons, and *s* is the external input with *s*_*E*_ and *s*_*I*_ denoting inputs to E and I neurons, respectively. *W* is the matrix of connection weights, including connections between E-to-E (*W*_*EE*_), E-to-I (*W*_*IE*_), I-to-E (*W*_*EI*_), and I-to-I (*W*_*II*_) neurons. *τ* is the effective time constant of the network integration, and []_+_ denotes half-wave rectification. We used forward Euler method to solve for the firing rates of neurons.

Spiking networks were modelled by simulating the equations describing the membrane potential dynamics of leaky integrate-and-fire neurons:

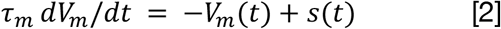

where *V*_*m*_ is the membrane potential of a neuron, and τ_m_ = *RC* is the time constant of integration of the membrane potential, with *R* and *C* denoting the membrane resistance and capacitance, respectively. When the membrane potential reaches a voltage threshold (*V*_*th*_), a spike is elicited and the membrane potential is reset to the reset voltage, *V*_*reset*_ = 0· *s*(*t*) = *R I*(*t*) describes the momentary input to the neurons, which arises from incoming spikes and comprises external (feedforward and non-local) input and recurrent input from presynaptic neurons in the network. Once a spike is emitted in a presynaptic source, an instantaneous change in the membrane potential of all postsynaptic sources is emulated in the next simulation time step, by the value of *w*, which is expressed in units of volts and describes the effect of *RI* simultaneously. The total input at time *t* for a postsynaptic neuron *i* is given by *s*(*t*) = ∑_*j*_ w_*ij*_ γ_*j*_ (*t*), where γ_*j*_ (*t*) denotes the presence (1) or absence (0) of spike in presynaptic sources, with w_*ij*_ describing the weight of connection from the *j*-th presynaptic source. We used exact integration method (*83*) to solve for the membrane potential and spiking activity of neurons.

Network connectivity is described by the weight matrix *W*, with *w*_*ij*_ denoting an entry on its *i*-th row and *j*-th column. Connections between neurons are established by drawing from a binomial distribution with probability ϵ (ϵ = 1 for networks with all-to-all connectivity). *w*_*ij*_ is set to zero if there is no connection from a pre- to postsynaptic neuron. If there is a connection, *w*_*ij*_ is drawn from a uniform distribution (with mean *J*) in randomly connected networks. In networks with specific connectivity, *w*_*ij*_ depends on the functional similarity of pre- and postsynaptic neurons. Neurons are assumed to have a 1D receptive field (e.g., orientation selectivity), and the weight of connections is modulated as:

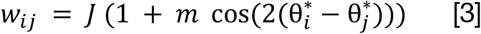

where *θ*_*i*_^*^ and *θ*_*j*_^*^ are the preferred orientation (in radians) of pre- and postsynaptic neurons, and *m* controls the degree of specificity of the connections (with *m* = 0 recapitulating random, nonspecific connectivity). Default parameters of simulations are listed in **Table 1**.

**Table 1.** Table of parameters.

| Description | Type | Param. | Fig. 1 | Figs. 2,3 | Figs. 4,5 | Fig. 6 | Fig. 7 |
| --- | --- | --- | --- | --- | --- | --- | --- |
| No. of neurons | E | $N_E$ | 500 | 500 | 400 | 200 | 400 |
| | I | $N_I$ | 500 | 500 | 400 | 200 | 400 |
| Time constant of neuronal integration | E&I | $\tau$ | 10 | 10 | 10 | 10 | 10 |
| Average weight of synaptic connections | E→E | $w_{EE}$ | 0.004 | 0.004 | 0.005 | 0.005 | 0.005 |
| | E→I | $w_{IE}$ | 0.004 (k=1)<br>0.016 (k=4) | 0.004 (k=1)<br>0.016 (k=4) | 0.005 (k=1)<br>0.02 (k=4) | 0.005 (k=1)<br>0.02 (k=4) | 0.005 (k=1)<br>0.02 (k=4) |
| | I→E | $w_{EI}$ | -0.004 (k=1)<br>-0.016 (k=4) | -0.004 (k=1)<br>-0.016 (k=4) | -0.005 (k=1)<br>-0.02 (k=4) | -0.005 (k=1)<br>-0.02 (k=4) | -0.005 (k=1)<br>-0.02 (k=4) |
| | I→I | $w_{II}$ | -0.004 (k=1)<br>-0.016 (k=4) | -0.004 (k=1)<br>-0.016 (k=4) | -0.005 (k=1)<br>-0.02 (k=4) | -0.005 (k=1)<br>-0.02 (k=4) | -0.005 (k=1)<br>-0.02 (k=4) |
| Connection probability | All | $\epsilon$ | 1 | 1 | 1 | 1 | 1 |
| Synaptic specificity | All | m | 0.5 | 0 | 0 | 0 | 0 |
| No. of perturbed neurons | E | $N_p$ | 10, 50, 100,<br>150, 200 | 10, 50, 100,<br>150, 200 | 20 | 20 | 100 |
| Time of pulse On/Off | | $T_p =$<br>$T_{on} = T_{off}$ | 10, 20, 50,<br>100 | 10, 20, 50,<br>100 | 50 | 100 | 50 |
| Stimulus (baseline) | E | $s$ | 1 | 1 | 1 | 1 | 1 |
| Perturbation size | E | $\delta s$ | 0.1 (k=1)<br>0.1 (k=4) | 0.1 (k=1)<br>0.1 (k=4) | 0.1 (k=1)<br>0.1 (k=4) | 0.1 (k=1)<br>0.1 (k=4) | 0.1 (k=1)<br>0.1 (k=4) |
| Learning rate | E→E | $\eta_{EE}$ | - | - | 0.2 | 0.1 | 0.1 |
| | E→I | $\eta_{IE}$ | - | - | 0 | 0 | 0.1 |
| | I→E | $\eta_{EI}$ | - | - | 0 | 0 | 0.1 |
| | I→I | $\eta_{II}$ | - | - | 0 | 0 | 0 |

### Network plasticity

To induce neuronal assemblies, a subset of *N*_*p*_ excitatory neurons in the network are perturbed. The perturbation pattern consists of *n*_*s*_ alternating pulses (ON/OFF); each pulse stays ON (*s*_*ON*_ = *s*_0_ + δ*s*) for *T*_*ON*_ and turns off (*s*_*OFF*_ = *s*_0_) for *T*_*OFF*_· *s*_0_ describes the input to the neurons before perturbations, respectively, and δ*s* denotes the strength of perturbation (e.g. corresponding to laser intensity in optogenetic stimulations (*19, 67*)). The total duration of perturbation is therefore *n* (*T*_*ON*_ + *T*_*OFF*_), with the duty cycle of *T*_*ON*_/(*T*_*ON*_ + *T*_*OFF*_). Assuming *T*_*p*_ = *T*_*ON*_ = *T*_*OFF*_, the stimulation frequency is *f*_*p*_ = 1/*T*_*p*_.

Following perturbations, synaptic plasticity is assumed to change the initial weight matrix as a result of network activity. The change in the weight *w*_*ij*_ is given as a function of the activity of pre- and post-synaptic neurons:

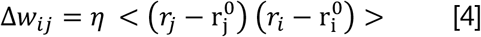

where *r*_*j*_ and *r*_*i*_ describe the firing rate of pre- and postsynaptic neurons, respectively, *η* is the learning rate, and <· > denotes the temporal average which is evaluated during perturbations. r^0^ denotes the average firing rate of individual neurons in their baseline state, obtained from network simulations before perturbations. We refer to this rule as covariance-based Hebbian learning, where covariance of the activity of pre- and postsynaptic neurons drives the plasticity. Two other versions of the rule are also considered, where response changes in only pre- or postsynaptic sources are considered, while the other term (post or pre, respectively) is still contributing to plasticity in absolute terms:

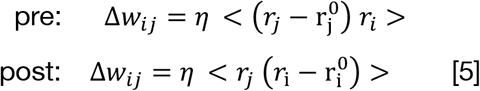

At each weight update, the weights of synapses are updated according to: *w*_*ij*_ ← *w*_*ij*_ + Δ *w*_*ij*_.

### Data analysis

To quantify the strength of assembly formation as a result of perturbation-induced plasticity we calculated ensemble potentiation. For each postsynaptic neuron (*i*) in the targeted pool of neurons (Ω), the sum of weight changes from the presynaptic sources within the pool (*j* ∈ Ω) is calculated: ∑_j∈Ω_ Δ *w*_*ij*_. Ensemble potentiation within the assembly (*E*_*w*_) is then obtained as the average of this value over postsynaptic neurons within the pool (i ∈ Ω):

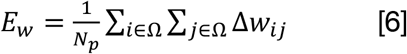

Out-of-assembly potentiation is, in turn, quantified by the average (over postsynaptic neurons within the targeted pool: i ∈ Ω) of the sum of presynaptic weight changes from excitatory neurons outside the assembly (*j* ∉ Ω):

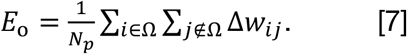

Specificity of induction is quantified by comparing within-assembly and out-of-assembly ensemble potentiation: (*E*_*w*_–*E*_*o*_)/(*E*_*w*_ + *E*_*o*_). To quantify the average weight changes within ensembles per individual synapse (e.g. as in **Fig. 2**, and as used in **Theoretical analysis** below), we also calculated average potentiation as: 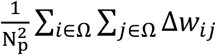.

Behavioral performance of induced neuronal assemblies was quantified by two metrics (**Fig. 6**). First, recall strength was used to measure the absolute capacity of neuronal ensembles to trigger readout responses upon partial stimulation. The activity of the linear readout was calculated as the sum (over neurons) of the activity of non-activated (NA) cells in the ensemble: *r*_*ro*_ = ∑_*i*∈*NA*_ *r*_*i*_. The average (temporal) differential response of the readout after partial activation was taken as a measure of recall strength: 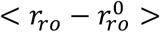, where 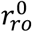 is the baseline activity of the readout and <· > denotes temporal averaging evaluated during partial activation.

A discriminability index was also developed to characterize how neuronal ensembles can distinguish between different behavioral contexts (A and B in **Fig. 6**). Using signal detection theory, it was calculated as:

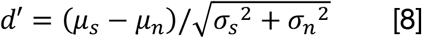

where µ_*s*_ and µ_*n*_ are the average readout responses to “signal” (relevant context) and “noise” (irrelevant context), respectively. µ_*s*_ is calculated as the average (across repetitions) of < *r*_*ro*_ >, when a small number of neurons in the same ensemble are activated, while µ_*n*_ corresponds to the condition where same number of neurons from the other ensemble are stimulated. σ is the std of < *r*_*ro*_ > over different repetitions of the partial stimulation in respective conditions.

### Theoretical analysis

To obtain theoretical insights into our numerical simulations, we analyzed how ensembles form in different E/I regimes and by different perturbation patterns. We first calculated the average potentiation of synapses expected from the linearized dynamics of the network. Writing Eq. 1 for the stationary state of network responses (*dr*/*dt* = 0), we have:

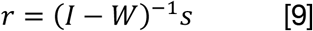

where 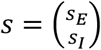 and 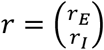 denote the *N* × 1 vectors of input and output activity, respectively (with *N* denoting the total number of E and I neurons in the network, *N* = *N*_*E*_ + *N*_*I*_). Perturbation of a subset of excitatory neurons by 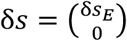 changes the output firing rates:

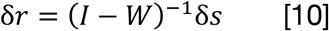

Applying the covariance-based Hebbian rule in Eq. 4, weight changes can be written as:

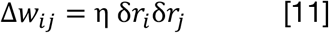

Matrix of weight changes *P*, with entry *p*_*ij*_ = Δ*w*_*ij*_ on *i*-th row and *j*-th column representing the weight change of the connection from the *i*-th presynaptic neuron to the *j*-th postsynaptic one, can, therefore, be expressed as:

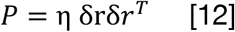

where S = δ*r*δ*r*^*T*^ is the covariance matrix of response changes following perturbations. Writing *A* = (*I* − *W*)^−1^ and substituting Eq. 10, matrix of plasticity *P* can in turn be expressed in terms of input perturbations as:

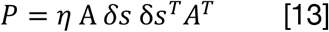

For different patterns of perturbations of excitatory neurons with different *N*_*p*_ and strength of perturbation, we can evaluate the ensemble potentiation from Eq. 12 by knowing the initial weight matrix, *W* (plotted in **Fig. 2** as prediction from theory based on W). Note that the prediction of this analysis by definition does not depend on *T*_*p*_, as it is based on stationary state responses. While the previous analysis sheds light on the relation between dynamics of the networks and the resulting weight changes via the weight matrix, it still needs to be evaluated numerically; especially, calculating *A* = (*I* − *W*)^-1^ can be computationally expensive for large matrices and precludes further analytical insights into the key parameters underlying the emergence of different regimes of induction. We therefore developed a mean-field analysis to calculate average potentiation within the assembly as a function of the average parameters of connectivity. The perturbed excitatory (*E*_1_), unperturbed excitatory (*E*_2_) and inhibitory (I) populations were reduced to single nodes in the mean-field analysis, with the connectivity between them described by:

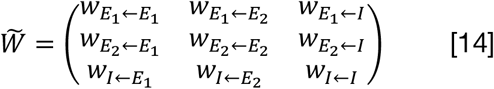

For the connectivity matrix parameterized in **Fig. 1G**, and assuming that a fraction *f* of E neurons are perturbed (*f* = *N*_*p*_ /N_E_), we can write the mean-field weight matrix as:

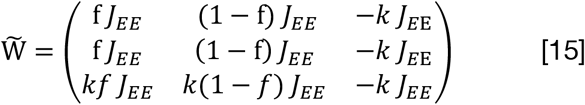

where *J*_*EE*_ is the mean-field, overall coupling strength of E population. For a network with connection probability ϵ and average weight *J*_5_ of individual E→E synapses, it can be expressed as *J*_*EE*_ = *N*_*E*_ ϵ*J*_0_. The mean-field coupling of I→{E,I} population can in turn be expressed as *N*_*I*_ϵ*gJ*_0_, where *g* determines the inhibition-dominance of individual I→{E,I} synapses over E→E ones. If *N*_*E*_ = *N*_*I*_, and given similar connection probabilities for all connection types, *k* = *g*. Dominant individual E→I synapses by the same factor also leads to overall dominance of E→I coupling: *N*_*E*_ 𝜖*kJ*_0_, as expressed in mean-field couplings in Eq. 15.

Knowing the mean-field matrix 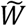, we can now obtain the corresponding matrix of weight changes for the mean-field analysis, 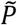, from Eq. 13, as:

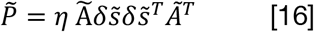

where 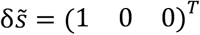 and 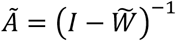. To obtain the average potentiation of synapses within the ensemble of perturbed neurons, we are interested in entry 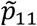 of the 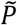 matrix, which can be obtained as:

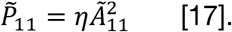

Writing *J*_*EE*_, Ã_11_ can in turn be computed from 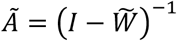 as:

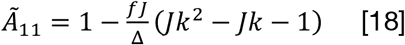

where Δ = *J*^2^ *k*^2^ − *J*^2^*k* + *Jk* − *J* + 1. This is used in **Fig. 2** for the mean-field analysis. For *k* = 1, Eq. 18 suggests that *Ã*_11_ ≈ 1 + *fJ*, and hence:

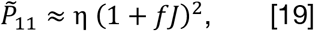

implying a supralinear enhancement of assembly formation for larger fraction of perturbed neurons. Note that this is the case independent of how large or small *J* is; especially whether *J* < 1 or *J* > 1 (unstable E-E subnetwork) does not change the result (cf. **Fig. 2C**). For large *J*, and strong *k* (*Jk* ≫ 1), another regime is obtained with Ã_11_ ≈ 1 − *f*, and

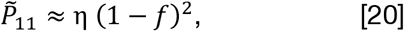

which suggests a weaker potentiation of synapses for larger *f* and hence *N*_*p*_ (cf. **Fig. 2F**).

## Acknowledgments

We thank Simon Peron and Ravi Pancholi for their comments on the manuscript, and Julia Gallinaro, Douglas Feitosa Tome and other member of the Clopath Lab for insightful discussions. This work was supported by BBSRC BB/N013956/1, BB/N019008/1, Wellcome Trust 200790/Z/16/Z, Simons Foundation 564408 and EPSRC EP/R035806/1. S.S and C.C designed the study; S.S performed the research; S.S and C.C wrote the paper. The authors declare that they have no competing interests. All data needed to evaluate the conclusions in the paper are present in the paper and/or the Supplementary Materials. Codes for reproducing main simulations are freely available from ModelDB (http://modeldb.yale.edu/266954).

## Supplementary figures

**Fig. S1.**
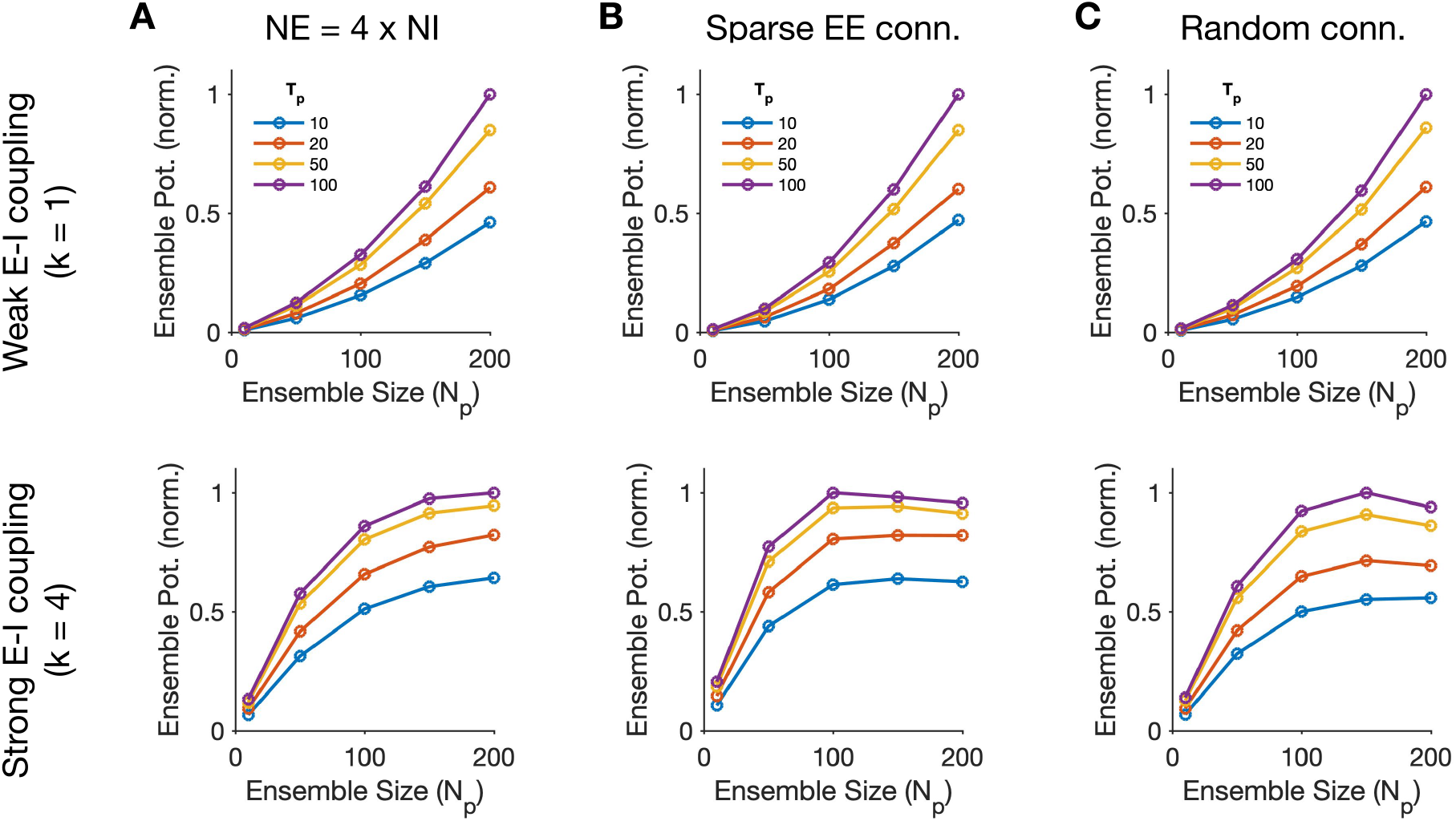
Induction of neuronal assemblies in networks with different combinations of parameters. **(A)** Same as **Fig. 1J,M**, respectively, for a network with larger number of excitatory neurons. *N*_*E*_ = 800, *N*_*I*_ =200. To provide the same level of inhibition, I→{E,I} weights are made 4 times stronger to adjust for the lower number of inhibitory neurons. (**B**) Same as **Fig. 1J,M**, for a network with sparser E-E connectivity. As opposed to all-to-all connectivity we assumed before, excitatory neurons are now connected to each other with 25% probability of connections (ϵ=0.25). To have the same level of overall E-E coupling, E→E weights are made 4 times stronger to adjust for the lower number of synapses. (**C**) Same as **Fig. 1J,M**, for a network with random connectivity. As opposed to network in **Fig. 1**, the weights of connections here are not modulated by functional similarity of neurons (see **Methods** for details).

**Fig. S2:**
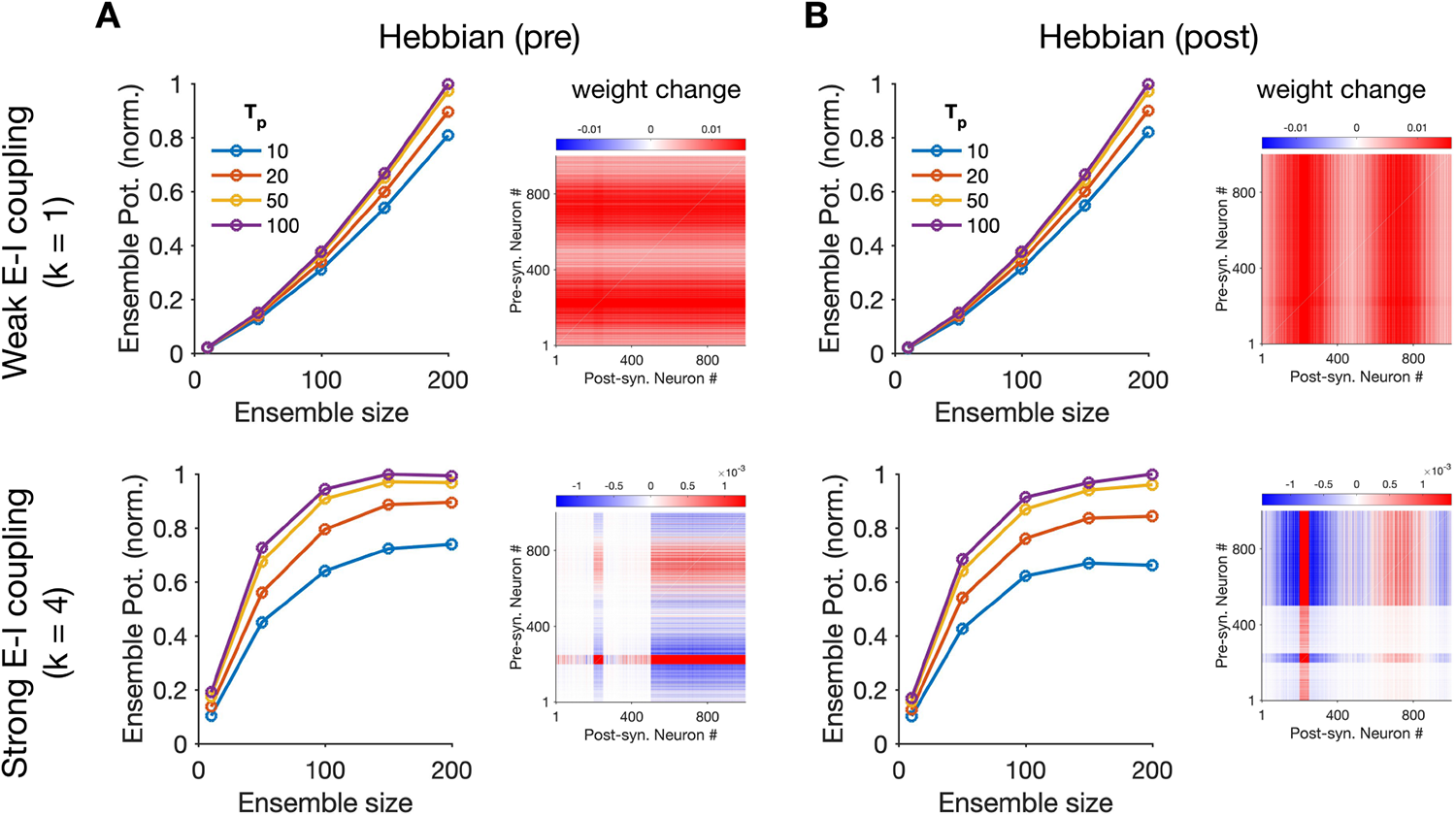
Induction of neuronal assemblies in networks with different Hebbian rules. The covariance-based Hebbian rule in **Fig. 1**, Δ*w*∝(*r*_*pre*_−*r*^0^_*pre*_)(*r*_*post*_−*r*^0^_*post*_) (see **Methods**), was governed by the deviation of pre- and post-synaptic activity from their baseline value before perturbations (denoted by *r*^0^). This is changed here to consider different Hebbian-type rules of plasticity which depend only on the deviations of pre-(A) or post-synaptic (B) activity, while the absolute value of post- or pre-synaptic activity is preserved, respectively. (**A**) Same as **Fig. 1J,M** when weight changes are governed by: 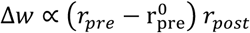. (**B**) Same as **Fig. 1J,M** when weight changes are governed by: Δ*w*∝*r*_*pre*_(*r*_*post*_−*r*^0^_*post*_). The matrix of weight changes for *N*_*p*_= 50 and *T*_*p*_ = 50 is shown for each condition.

**Fig. S3:**
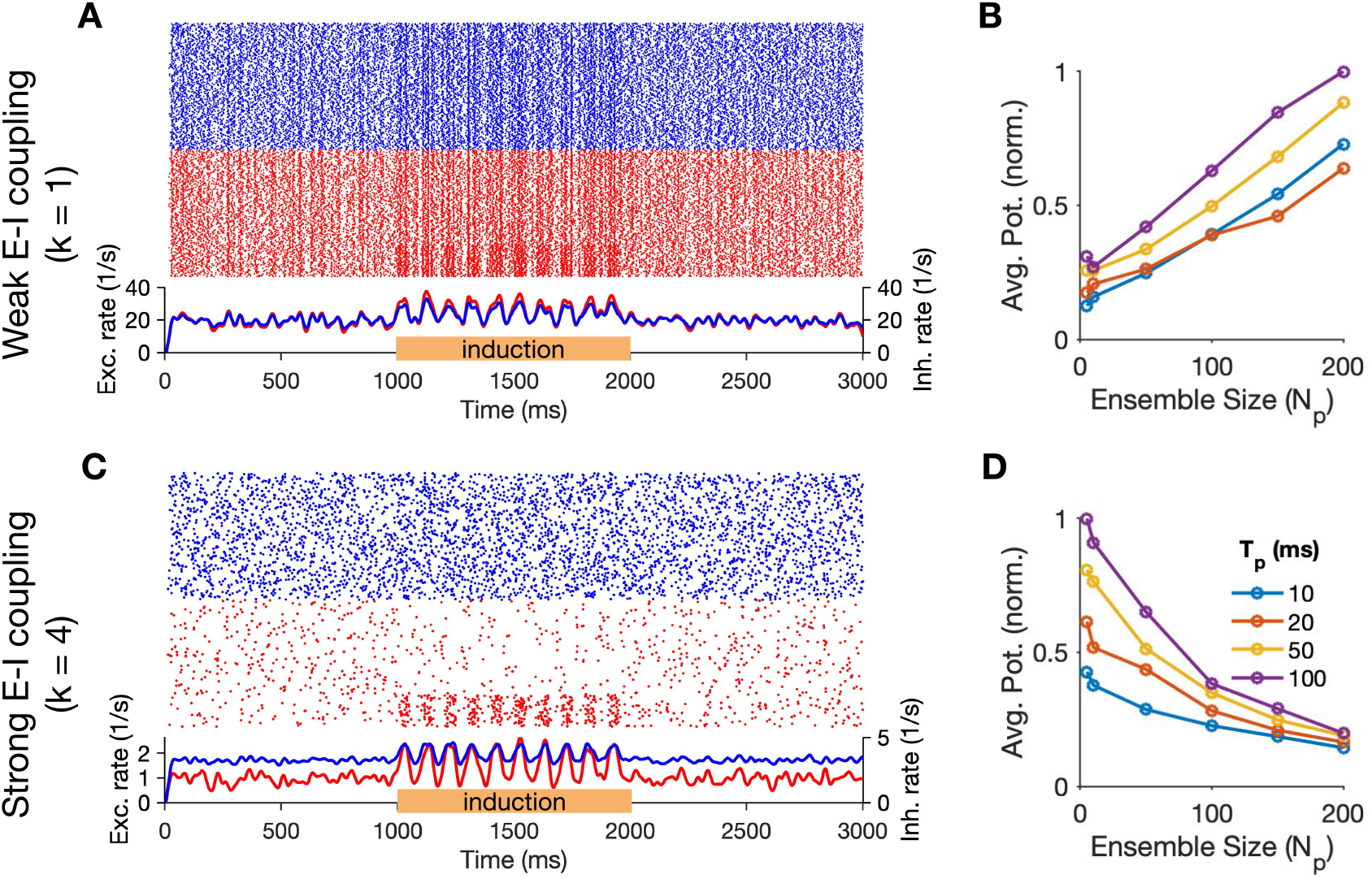
Induction of neuronal assemblies in spiking networks. (**A**) Induction of neuronal assemblies in networks of spiking neurons in the weak E-I coupling regime (*k* = 1). A sample simulation (with *N*_*p*_= 100 and *T* = 50 *ms*) is shown, with raster plots of activity on top (red: Exc., blue: Inh.), and average population activity of *N*_*E*_ Exc. and *N*_*I*_ Inh. neurons on the bottom (calculated in bins of 1 ms and smoothened with a sliding Gaussian kernel of 20 ms length).*N*_*E*_ = *N*_*I*_ = 400. Connectivity is all-to-all (ϵ= 1) and random (*m* = 0), with an average weight of *w*_*EE*_ = 0.1 *mV* for E-to-E connections. τ^*E*^_*m*_ = τ^*I*^_*m*_ = 20 ms. Perturbations are performed for 10 cycles in this example. (**B**) Average potentiation of individual synapses for induction protocols with different values of *N*_*P*_ and *T*_*P*_ shows similar dependence on the size of induced ensembles as rate-based models (cf. **Fig. 2A**). The plasticity of synapses is governed by a Hebbian plasticity rule based on the covariances of pre- and post-synaptic sources. The pre-term is read from the presynaptic spiking activity of neurons and the post-term is inferred from the average free membrane potential of postsynaptic neurons (see **Methods** for details). The absolute value of the average potentiation of synapses within induced assemblies are normalized to the maximum value across all inductions. Perturbations are performed for 100 cycles to obtain more reliable estimates of response changes. (**C,D**) Same as (A,B) for induction in spiking networks with *k* = 4.

**Fig. S4:**
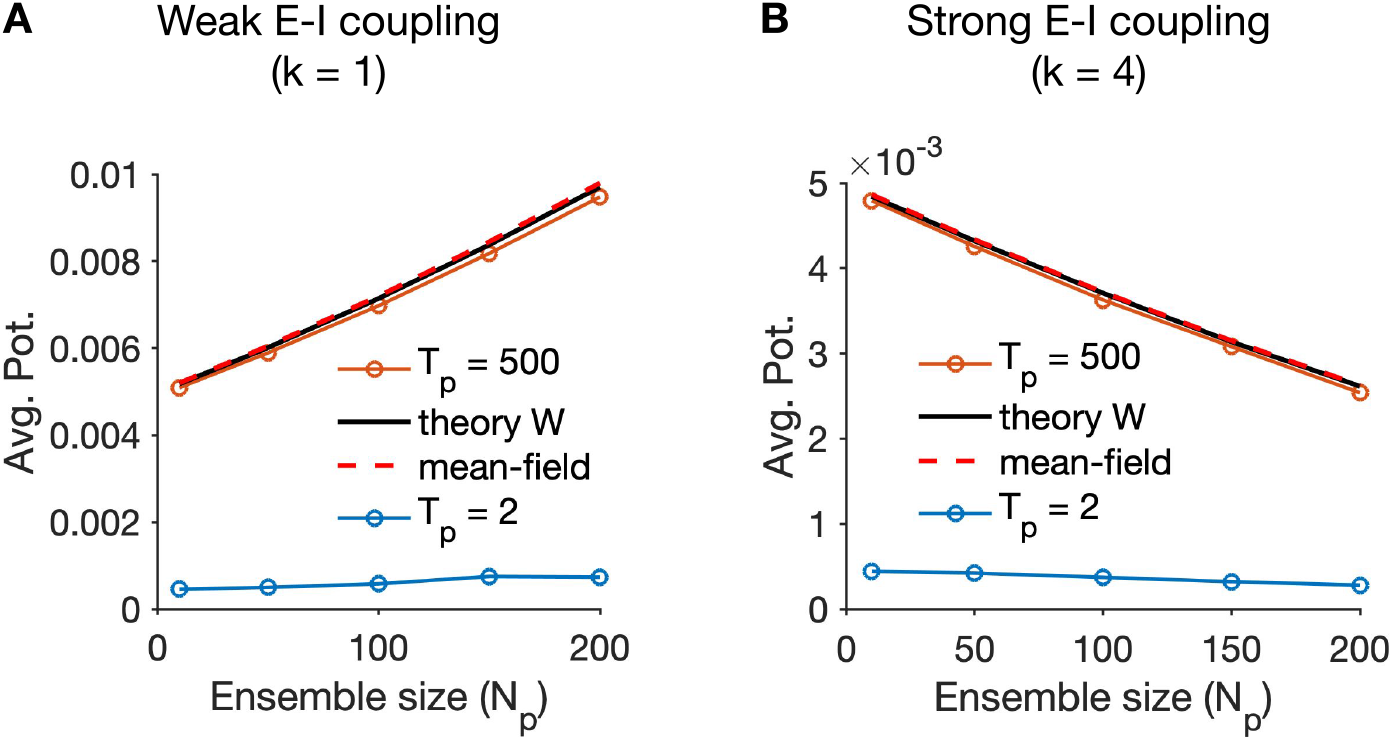
Potentiation of synapses with extreme perturbation times. (**A,B**) Same as **Fig. 2B,E**, respectively, for extremely small and large values of *T*_*p*_. The residual discrepancy between theory and simulations (cf. **Fig. 2B,E**) is absent for very long pulses (*T*_*p*_= 500), while the increasing/decreasing trends with ensemble size are much weaker for very brief perturbations (*T*_*p*_= 2).

**Fig. S5:**
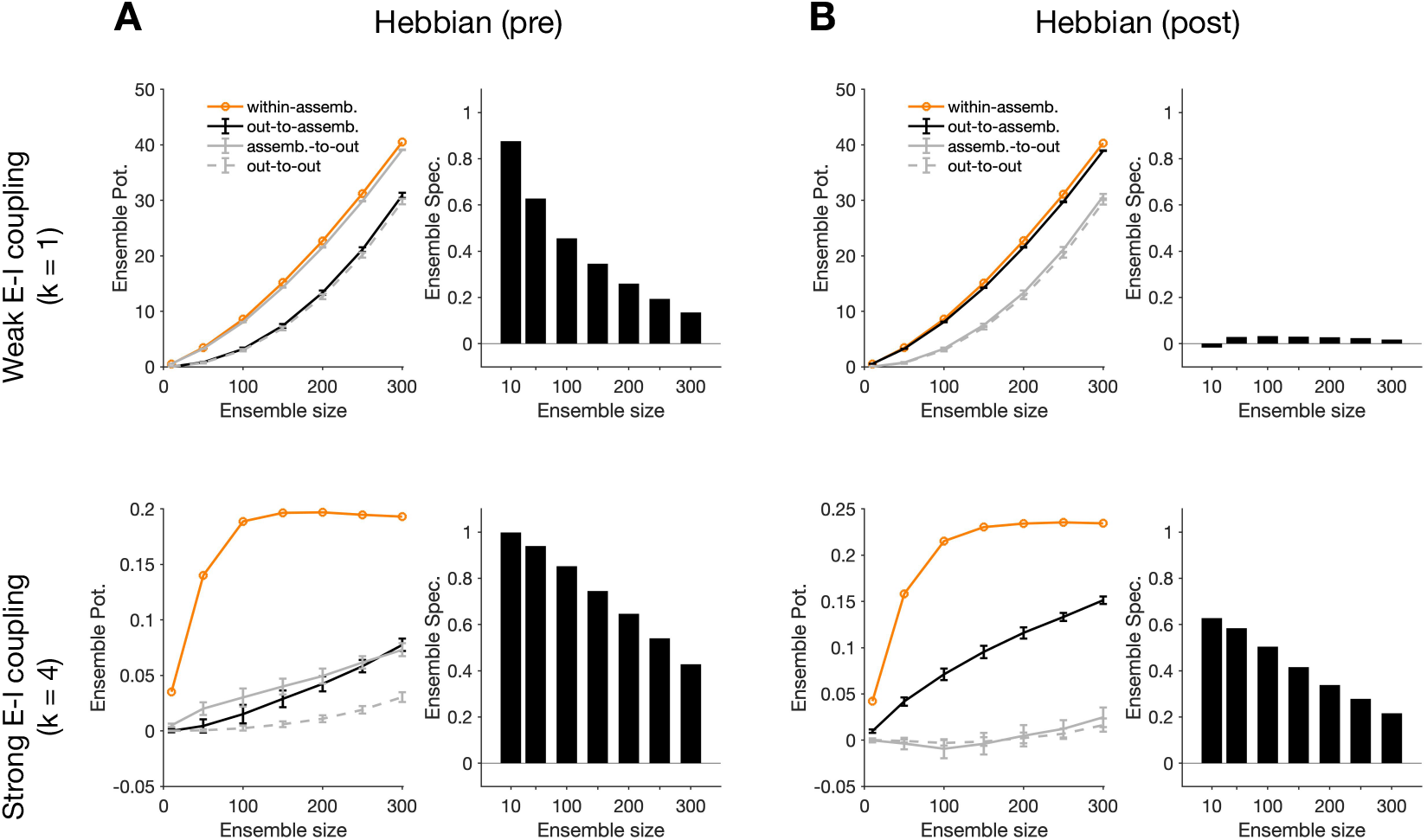
Specificity of assembly formation with different Hebbian rules. (**A,B**) Specificity of ensemble formations (similar to **Fig. 3B-E**) for different Hebbian rules of plasticity. As opposed to the covariance-based rule in **Fig. 1** which depended on the response changes of both pre- and post-synaptic neurons (see **Methods**), here the Hebbian rules depend on pre-(A) or post-synaptic (B) changes, while preserving the dependence on the absolute activity of the post and pre, respectively (similar to rules in **Fig. S2A,B**, respectively; see the caption for the details of the rules employed here).

**Fig. S6:**
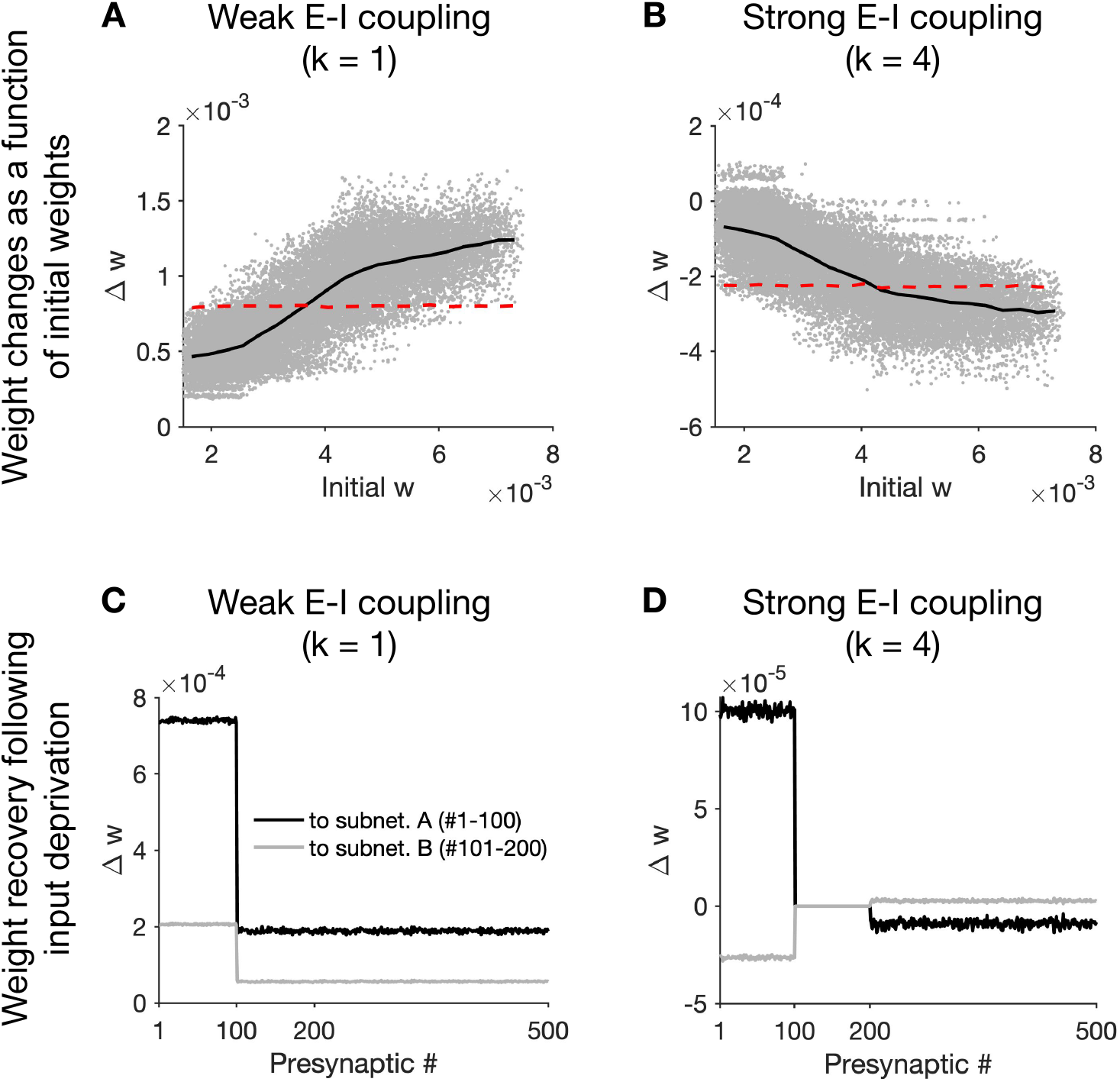
Specificity of ensemble potentiation based on initial weights and after input deprivation. (**A,B**) Weight changes (Δ*w*) of out-of-assembly connections (same as **Fig. 1I,L**, respectively), for different regimes of E-I coupling (A and B, respectively), as a function of their initial connectivity (Initial w) to the ensemble. The initial ensemble, namely the E neurons perturbed initially, are *N*_*p*_= 50 neurons with similar preferred orientations (cf. **Fig. 1I,L**). The black lines show the average value of Δ*w* calculated in 20 equal bins along the x-axis, respectively. The red lines show similar average values of Δ*w*, when *N*_*p*_= 50 neurons in the initial ensemble are chosen randomly, independent of their initial preferred orientations. (**C,D**) Weight changes of subnetworks with deprived input. In networks similar to those in **Fig. 2** (with random connectivity), the feedforward input to a fraction of neurons (#1-200) is reduced to half the initial value. Correlated input patterns (similar to those delivered in **Fig. 1**, with *T*_*p*_= 50) from other sources are assumed to activate one of the subnetworks (A: #1-100), while the other subnetwork (B: #101-200) does not receive the input perturbations. The average weight changes of presynaptic E neurons (*N*_*E*_ = 500) to different subnetworks (cf. e.g. **Fig. 2**) are plotted, for weak (C) and strong (D) E-I coupling regimes, respectively.

**Fig. S7:**
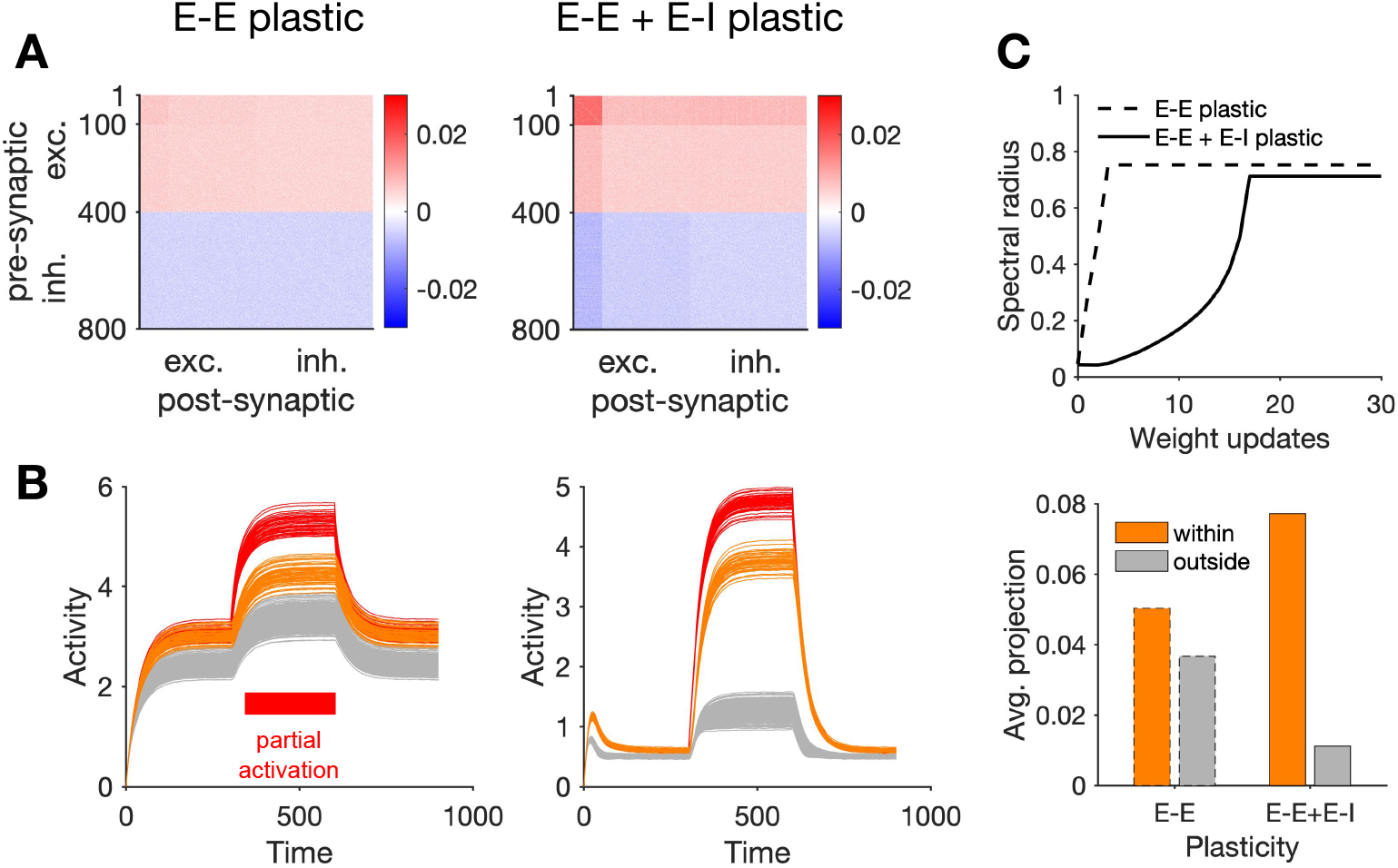
Transition between different regimes of induction with different rules of E-I learning. Same as **Fig. 7** for a different rule of E-I plasticity. E→I and I→E plasticity in **Fig. 7** depended on changes in the activity of pre-synaptic excitatory and inhibitory neurons, respectively (**Methods**). Here, pre- and post-synaptic activity changes are both considered for both types of synapses (see **Methods** for details).

